# Brain-wide neuronal circuit connectome of human glioblastoma

**DOI:** 10.1101/2024.03.01.583047

**Authors:** Yusha Sun, Xin Wang, Daniel Y. Zhang, Zhijian Zhang, Janardhan P. Bhattarai, Yingqi Wang, Weifan Dong, Feng Zhang, Kristen H. Park, Jamie Galanaugh, Abhijeet Sambangi, Qian Yang, Sang Hoon Kim, Garrett Wheeler, Tiago Gonçalves, Qing Wang, Daniel Geschwind, Riki Kawaguchi, Huadong Wang, Fuqiang Xu, Zev A. Binder, H. Isaac Chen, Emily Ling-Lin Pai, Sara Stone, MacLean P. Nasrallah, Kimberly M. Christian, Marc Fuccillo, Donald M. O’Rourke, Minghong Ma, Guo-li Ming, Hongjun Song

**Affiliations:** Neuroscience Graduate Group, Perelman School of Medicine, University of Pennsylvania, Philadelphia, PA, USA; Department of Neuroscience and Mahoney Institute for Neurosciences, Perelman School of Medicine, University of Pennsylvania, Philadelphia, PA, USA; Department of Neurosurgery, Perelman School of Medicine, University of Pennsylvania, Philadelphia, PA, USA; Sidney Kimmel Medical College, Thomas Jefferson University Hospital, Philadelphia, PA, USA; Department of Neuroscience and Gottesman Institute for Stem Cell Biology and Regenerative Medicine, Albert Einstein College of Medicine, Bronx, NY, USA; Program in Neurogenetics, Department of Neurology, David Geffen School of Medicine, University of California Los Angeles, Los Angeles, CA, USA; Shenzhen Key Laboratory of Viral Vectors for Biomedicine, Shenzhen-Hong Kong Institute of Brain Science, Shenzhen Institute of Advanced Technology, Chinese Academy of Sciences, Shenzhen, China; Glioblastoma Translational Center of Excellence, The Abramson Cancer Center, University of Pennsylvania, Philadelphia, PA, USA; Institute for Regenerative Medicine, University of Pennsylvania, Philadelphia, PA, USA; Corporal Michael J. Crescenz Veterans Affairs Medical Center, Philadelphia PA, USA; Department of Pathology and Laboratory Medicine, University of Pennsylvania, Philadelphia, PA, USA; Department of Cell and Developmental Biology, Perelman School of Medicine, University of Pennsylvania, Philadelphia, PA, USA; Department of Psychiatry, Perelman School of Medicine, University of Pennsylvania, Philadelphia, PA, USA; The Epigenetics Institute, Perelman School of Medicine, University of Pennsylvania, Philadelphia, PA, USA

**Author notes:** Corresponding authors (G.M.) and (H.S.). These authors contributed equally to this work (Yusha Sun, Xin Wang).

## Abstract

Glioblastoma (GBM), a universally fatal brain cancer, infiltrates the brain and can be synaptically innervated by neurons, which drives tumor progression^1–6^. Synaptic inputs onto GBM cells identified so far are largely short-range and glutamatergic^7–9^. The extent of integration of GBM cells into brain-wide neuronal circuitry is not well understood. Here we applied a rabies virus-mediated retrograde monosynaptic tracing approach^10–12^ to systematically investigate circuit integration of human GBM organoids transplanted into adult mice. We found that GBM cells from multiple patients rapidly integrated into brain-wide neuronal circuits and exhibited diverse local and long-range connectivity. Beyond glutamatergic inputs, we identified a variety of neuromodulatory inputs across the brain, including cholinergic inputs from the basal forebrain. Acute acetylcholine stimulation induced sustained calcium oscillations and long-lasting transcriptional reprogramming of GBM cells into a more invasive state via the metabotropic CHRM3 receptor. CHRM3 downregulation suppressed GBM cell invasion, proliferation, and survival *in vitro* and *in vivo*. Together, these results reveal the capacity of human GBM cells to rapidly and robustly integrate into anatomically and molecularly diverse neuronal circuitry in the adult brain and support a model wherein rapid synapse formation onto GBM cells and transient activation of upstream neurons may lead to a long-lasting increase in fitness to promote tumor infiltration and progression.

## Main Text

GBM is the most common and deadly primary brain cancer in adults and is characterized by its heterogeneity^13^, complex tumor microenvironment^14^, and invasiveness^15^. A growing body of evidence in the emerging field of cancer neuroscience suggests that circuit integration of glioma drives tumor progression, invasion, and decreased patient survival^7,9,15–20^. Given that GBM cells are highly infiltrative, synapses onto these migratory cells will inevitably be transient. Whether transient synapses can exert long-lasting influences on the behavior of migratory GBM cells is still unclear. Additionally, synaptic inputs onto glioma cells identified so far have been limited to local glutamatergic projections^7,8^, and the circuit architecture and neuronal subtype diversity of neuron-glioma synaptic connections remain to be elucidated. Monosynaptically-restricted transsynaptic tracing using modified rabies virus is a classic methodology in neuroscience to systematically map synaptic inputs onto defined targets, or starter cells^21^. This strategy has been widely employed to elucidate whole-brain neuronal networks with specific neurons^22,23^ or oligodendrocyte precursor cells^24^ as starter cells. Here we performed *in vivo* retrograde monosynaptic rabies virus tracing of transplanted patient-derived GBM organoids (GBOs)^25,26^ to characterize the landscape of neuronal innervation onto GBM cells and further investigated functional effects of neuromodulatory inputs onto GBM cells.

### Expression of diverse neurotransmitter receptors in GBM

To explore the potential for GBM cells to respond to different neurotransmitters, we used single-cell RNA sequencing (scRNAseq) via a Smart-seq3-based protocol^27^ for deep characterization of neurotransmitter receptor expression (*n* = ∼6,200 genes per cell). To account for significant intra- and inter-tumoral heterogeneity, we examined GBOs derived from genetically distinct, isocitrate dehydrogenase-wild type (IDH-wt) GBM tumors resected from three different patients (UP-9096, UP-9121, and UP-10072; Fig. 1a, Extended Data Fig. 1a-c). We found that human GBM cells expressed a wide variety of neurotransmitter receptors, including ionotropic and metabotropic glutamatergic, GABAergic, and cholinergic receptors as well as serotonergic, adrenergic, and dopaminergic receptors (Fig. 1a). Expression levels of these receptors were comparable to those in neural stem cells (NSCs) from human induced pluripotent stem cell (iPSC)-derived cortical organoids that we profiled in parallel^28^ (Fig. 1a, Extended Data Fig. 1d-e). We obtained similar results from our analysis of published scRNAseq datasets of adult primary IDH-wt GBM^13,29^ (Fig. 1a, Extended Data Fig. 1f). Consistent with prior studies^8^, GBM cells from all datasets also abundantly expressed post-synaptic scaffold genes, such as *HOMER1* and *DLG4* (encoding PSD-95) (Fig. 1a). Expression of neurotransmitter receptors and gene signature enrichment scoring for post-synaptic density genes revealed largely similar levels of expression among IDH-wt GBM cells across different cellular states^13^ in all datasets and in NSCs from organoids, with a slight enrichment in non-mesenchymal states and in peripheral infiltrating GBM cells compared to the tumor core in a primary tumor dataset^30^ (Extended Data Fig. 1g-j). These results reveal the capacity for GBM cells across different cellular states to receive and respond to synaptic inputs of diverse neurotransmitters.

**Figure 1.**
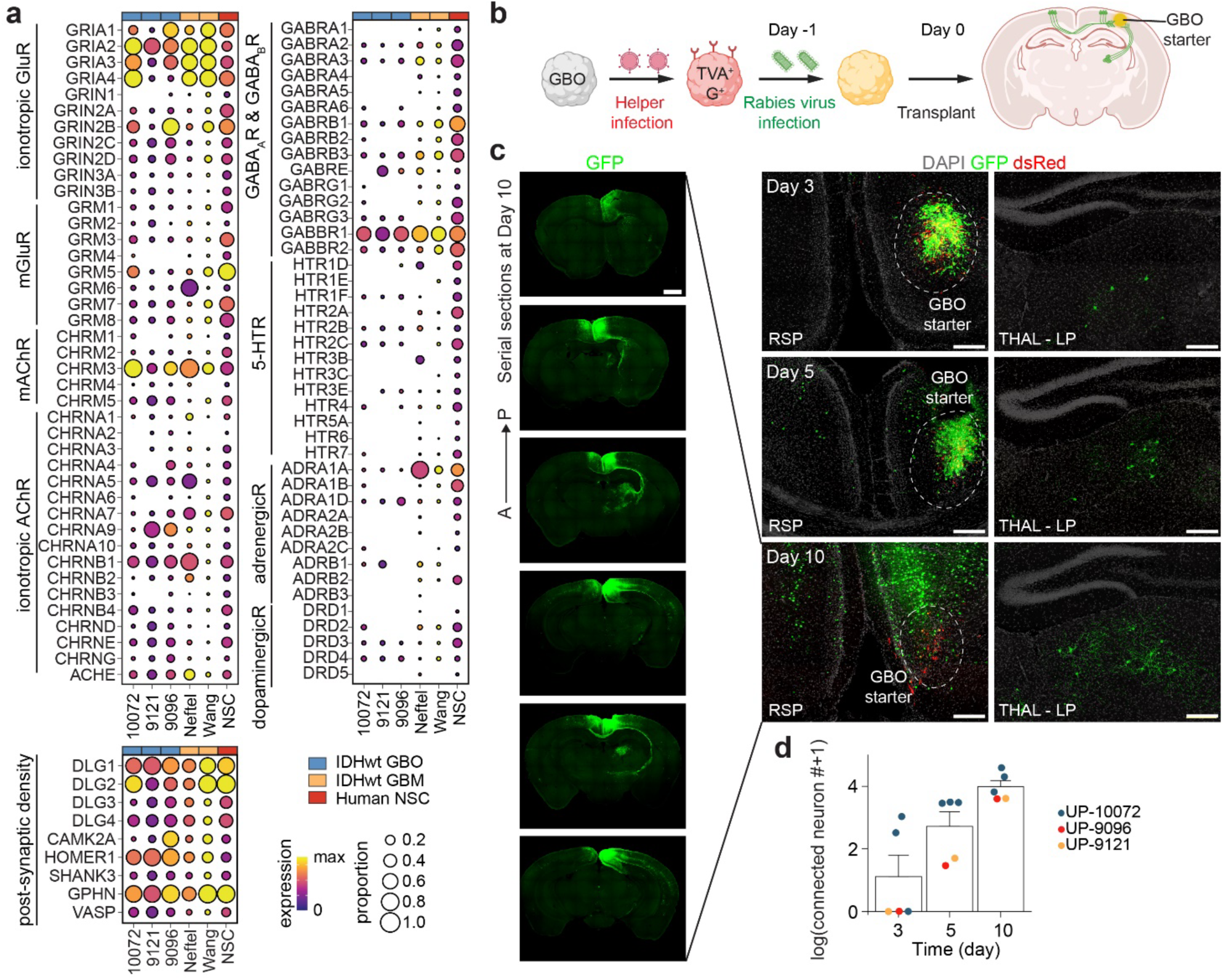
Rapid neuronal circuit integration of transplanted human GBM cells in the adult mouse brain. **a**, Expression of neurotransmitter receptor and post-synaptic density genes across single-cell transcriptomes of adult GBM (blue: UP-10072, UP-9121, UP-9096, *n* = 3 patient-derived GBOs; orange: Neftel et al.^13^, *n* = 20 patients, and Wang et al.^29^, *n =* 6 patients) and neural stem cells (NSCs) in 100-day old neocortical organoids (red: *n* = 3 organoids). Data are plotted as log-normalized counts, and the size of dots represents the proportion of cells in which the given gene is detected. **b**, Schematic illustrations of the transplantation paradigm of rabies virus pre-infected GBOs into immunodeficient mice. **c**, Sample confocal images of local (retrosplenial cortex, RSP) and long-range (lateral posterior thalamus, TH - LP) regions at 3, 5, and 10 days post transplantation (dpt) into the RSP. Scale bars, 200 μm. Left, representative serial sections from anterior to posterior at 10 dpt. Scale bar, 500 μm. See additional sample images in **Extended Data** Fig. 2d. **d**, Quantification of the number of labeled neurons at 3, 5, and 10 dpt. Each dot represents data from one mouse, and color represents GBOs from different patients. Values represent mean ± s.e.m. (*n* = 3 patients).

### Rapid neuronal circuit integration of GBM cells

To systematically map brain-wide neuronal projections onto GBOs after transplantation into the adult mouse brain, we employed monosynaptic rabies virus tracing^10–12^. Cultured GBOs were retrovirally transduced to express a DsRed reporter (R), the EnvA receptor TVA (T), and rabies virus glycoprotein (G) (named RTG; Extended Data Fig. 2a). Starter GBO cells directly infected by G protein-deleted EnvA-pseudotyped rabies virus expressing a GFP reporter (ΔG rabies virus)^31^ indicated by DsRed and GFP co-expression can retrogradely transmit rabies virus to their presynaptic neurons, which can be identified by GFP expression alone (Extended Data Fig. 2b). As these first-order presynaptic neurons do not express G protein, ΔG rabies virus is unable to further propagate, resulting in the monosynaptic nature of this tracing^31^ (Extended Data Fig. 2a-b). GBOs that were transduced with RTG retrovirus were efficiently infected by ΔG rabies virus, while control GBOs without RTG expression were not infected by ΔG rabies virus (Extended Data Fig. 2c).

Following a paradigm to map synaptic inputs onto transplanted brain organoids^23^, we pre-infected GBOs derived from three patients with ΔG rabies virus for orthotopic transplantation into the retrosplenial cortex (RSP) of adult immunodeficient mouse brains (Fig. 1b). We observed GFP expression in local (ipsilateral cortex, hippocampus) and long-range (ipsilateral thalamus, contralateral cortex, basal forebrain) projecting neurons beginning at 3 days post transplantation (dpt), with extensive labeling on the order of ∼10^3^ – 10^4^ neurons across GBOs derived from three patients by 10 dpt (Fig. 1c-d, Extended Data Fig. 2d). As it takes at least 2 days for rabies virus to replicate, retrogradely transmit across synapses, and sufficiently label cells^31^, the detection of GFP^+^ neurons as early as 3 dpt suggests strikingly rapid neuronal circuit integration of GBM cells *in vivo*.

We also performed control experiments to ensure the fidelity of our approach. First, we engineered a control helper retroviral construct by deleting the rabies virus glycoprotein coding sequence (RT), such that GBO starter cells transduced by RT retrovirus can be infected by ΔG rabies virus via TVA but cannot transmit it to upstream neurons (Extended Data Fig. 2e). As expected, upon transplantation, we detected DsRed^+^GFP^+^ starter GBM cells but no GFP^+^ mouse neurons (Extended Data Fig. 2f). Second, to rule out the possibility of non-specific labeling due to leakage of rabies virus from pre-infected GBM cells, we induced lysis of starter GBO cells to extract infection-competent rabies virus prior to transplantation. We found only very rare GFP^+^ neurons during the same time window (Extended Data Fig. 2g). Third, immunostaining for the glial marker GFAP did not reveal co-labeling with GFP either near or distant to the RSP injection site, consistent with the transsynaptic nature of rabies virus transmission^32^ (Extended Data Fig. 2h).

Together, this G protein-dependent monosynaptic rabies virus tracing system uncovered rapid and robust neuronal connectivity of transplanted human GBM cells in the adult mouse brain.

### Brain-wide anatomic atlas of synaptic inputs onto GBM cells

We next systematically characterized the brain-wide distribution of rabies virus-labeled neurons after transplantation of pre-infected GBOs from the three patients into four cortical and subcortical sites: primary somatosensory cortex (S1), primary motor cortex (M1), retrosplenial cortex (RSP), and hippocampus (HIP), which correspond to common anatomical regions where glioma appears in patients^33,34^ (Fig. 2a-d). At 10 dpt, we observed broadly distributed GFP^+^ cells throughout brain regions for GBOs from all three patients (Fig. 2a-e, Extended Data Fig. 3 and 4a), owing partially to the ability of these cells to rapidly infiltrate as indicated by the location of starter GBM cells (Fig. 2f). Transplantation of GBOs across different patients showed largely similar distributions of GFP^+^ cells for each transplantation site, suggesting conserved neuron-GBM interaction patterns despite the heterogeneity of GBM (Fig. 2e). Quantification of the proportion of GFP^+^ cells by brain region revealed that GBM cells in cortical areas received the highest proportion of inputs from the isocortex and secondarily from the thalamus (Fig. 2g). Cortical inputs onto GBM cells in both S1 and M1 were largely comprised of neurons in the sensory and motor cortex both ipsilaterally and contralaterally (Fig. 2a-b, e), reflecting the close functional association of these areas^35^. Contralateral neurons accounted for nearly 20% of total GFP^+^ cortical neurons (Fig. 2h), with L2/3 contralateral neurons as the dominant input subpopulation compared to those of L5 or L6 (Extended Data Fig. 4b), highlighting long-range cortical networks as substantial components of neuron-GBM circuitry^20^. Thalamic projections onto GBM cells, such as from ventral posteromedial and posterior complex thalamus upon S1 transplantation and from ventromedial thalamus upon M1/RSP transplantation (Fig. 2a-c, e), were almost entirely ipsilateral, consistent with known thalamo-cortical wiring^36,37^. For subcortical HIP transplantation, the most abundant inputs were found in the hippocampal (dentate gyrus, CA1, CA3) and retrohippocampal (subiculum, entorhinal cortex) areas (Fig. 2d, e, g). We also found GFP^+^ neurons in diverse subcortical regions, including the hypothalamus, claustrum, and midbrain, for all transplanted sites, as well as the diagonal band nucleus (NDB) and medial septal nucleus (MS) of the basal forebrain upon RSP and HIP transplantations (Fig. 2c-e, Extended Data Fig. 4c-e).

**Figure 2.**
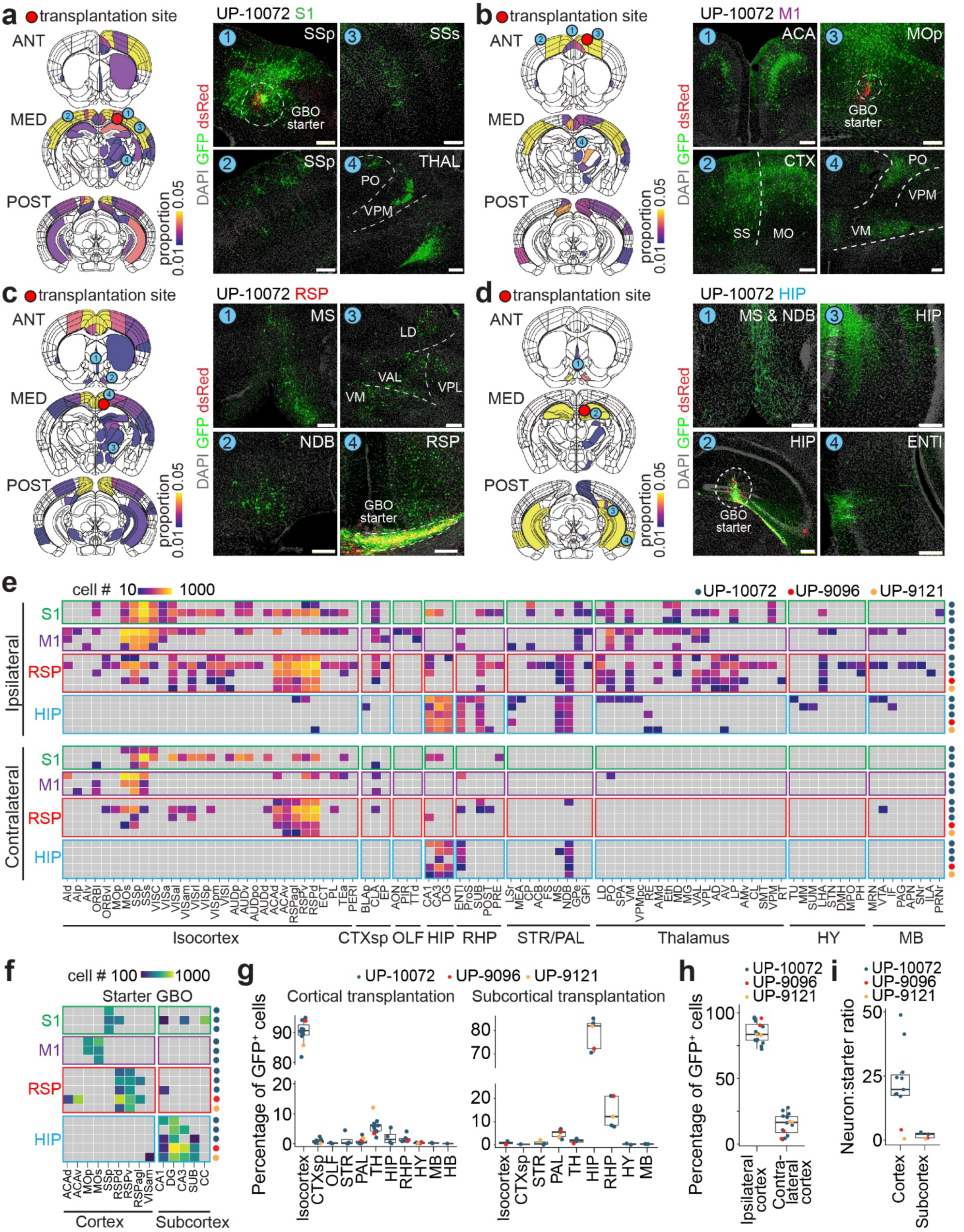
GBM cell input connectome with diverse anatomical projections. **a**, Left shows representative coronal brain section diagrams corresponding to anterior (ANT), medial (MED), and posterior (POST) adult mouse brain from the reference Allen Brain Mouse Atlas^99^ for primary somatosensory cortex (S1) transplantations. Areas are colored by normalized proportion of GFP^+^ neurons observed in corresponding brain regions relative to total GFP^+^ neuron number in the ipsilateral or contralateral hemispheres at 10 dpt (quantified from *n* ≥ 3 mice). Right shows sample confocal images of ipsilateral S1 (injection site), ipsilateral secondary somatosensory cortex (S2), contralateral S1, and ipsilateral thalamus with locations indicated on the slice diagrams on the left. Note that all images are oriented such that the right side of the image is the GBO transplantation site (ipsilateral). Starter GBM cells are circled with a dashed white line and labeled. THAL, thalamus; PO, posterior complex; VPM, ventral posteromedial. Scale bars, 200 μm. **b**, Same as **a** but for primary motor cortex (M1) transplantations (*n* = 3 mice). ACA, anterior cingulate area; CTX, cortex; VM, ventral medial. **c**, Same as **a** but for retrosplenial cortex (RSP) transplantations (*n* = 5 mice). MS, medial septum; NDB, diagonal band nucleus; VAL, ventrolateral; LD, laterodorsal; VPL, ventral posterolateral. **d**, Same as **a** but for hippocampal (HIP) area transplantations (*n* = 5 mice). ENTl, entorhinal. **e**, Heatmap of input projections to GBM cells, colored by GFP^+^ neuron number, grouped by transplantation location, and arranged by ipsilateral versus contralateral hemisphere. Columns in the heatmap represent individual brain regions as identified by the Allen Mouse Brain Atlas^99^ and are organized by larger brain regions. Each row represents an individual experiment at 10 dpt (*n* = 16 mice from GBOs derived from *n* = 3 patients). In this and subsequent Figures, GBO transplantation locations and patient ID are color-coded as indicated. CTXsp, cortical subplate; OLF, olfactory area; RHP, retrohippocampal area; STR, striatum; PAL, pallidum; HY, hypothalamus; MB, midbrain; AId, agranular insular area, dorsal part; AIp, agranular insular area, posterior part; AIv, agranular insular area, ventral part; ORBl, orbital area, lateral part; ORBvl, orbital area, ventrolateral part; MOp, primary motor area; MOs, secondary motor area; SSp, primary somatosensory area; SSs, secondary somatosensory area; VISC, visceral area; VISa, anterior visual area; VISal, anterolateral visual area; VISam, anteromedial visual area; VISrl, rostrolateral visual area; VISp, primary visual area; VISpm, posteromedial visual area; VISl, lateral visual area; AUDp, primary auditory area; AUDv, ventral auditory area; AUDpo, posterior auditory area; AUDd, dorsal auditory area; ACAd, anterior cingulate area, dorsal part; ACAv, anterior cingulate area, ventral part; RSPagl, retrosplenial area, lateral agranular part; RSPv, retrosplenial area, ventral part; RSPd, retrosplenial area, dorsal part; ECT, ectorhinal area; PL, prelimbic area; TEa, temporal association area; PERI, perirhinal area; BLAp, basolateral amygdalar nucleus, posterior part; CLA, claustrum; EP, endopiriform nucleus; AON, anterior olfactory nucleus; PIR, piriform area; TTd, taenia tecta, dorsal part; DG, dentate gyrus; ENTl, entorhinal area, lateral part; ProS, prosubiculum; SUB, subiculum; POST, postsubiculum; PRE, presubiculum; LSr, lateral septal nucleus, rostral part; MEA, medial amygdalar nucleus; CP, caudoputamen; ACB, nucleus accumbens; FS, fundus of striatum; MS, medial septal nucleus; NDB, diagonal band nucleus; GPe, globus pallidus, external segment; GPi, globus pallidus, internal segment; LD, lateral dorsal nucleus; PO, posterior complex; SPA, subparafascicular area; VM, ventral medial nucleus; VPMpc, ventral posteromedial, parvicellular part; RE, nucleus of reuniens; AMd, anteromedial nucleus, dorsal part; Eth, ethmoid nucleus; MD, mediodorsal nucleus; MG, medial geniculate complex; VAL, ventral anterior-lateral complex; VPL, ventral posterolateral nucleus; AD, anterodorsal nucleus; AV, anteroventral nucleus; LP, lateral dorsal nucleus; AMv, anteromedial nucleus, ventral part; CL, central lateral nucleus; SMT, submedial nucleus; VPM, ventral posteromedial nucleus; RT, reticular nucleus; TU, tuberal nucleus; MM, medial mamillary nucleus; SUM, supramamillary nucleus; LHA, lateral hypothalamic area; STN, subthalamic nucleus; DMH, dorsomedial nucleus; MPO, medial preoptic area; PH, posterior hypothalamic nucleus; MRN, midbrain reticular nucleus; VTA, ventral tegmental area; IF, interfascicular nucleus raphe; PAG, periaqueductal gray; APN, anterior pretectal nucleus; SNr, substantia nigra, reticular part; ILA, infralimbic area; PRNr, pontine reticular nucleus. **f**, Heatmap of starter GBM cell distribution with the same columnar labeling as in **e** for the same set of experiments, colored by GBO cell number and arranged by cortical versus subcortical regions. CC, corpus callosum. **g**, Proportions of input neurons across brain regions for subcortical (HIP, *n* = 5 mice) and cortical (S1, M1, and RSP, *n* = 11 mice) transplantation experiments at 10 dpt, colored by GBOs from different patients. Each dot represents data from one mouse. **h**, Proportion of input cortical neurons arising from either the ipsilateral or contralateral hemisphere for experiments as defined in **g**. **i**, Quantification of input neuron to starter GBM cell ratio for subcortical and cortical transplantation experiments as defined in **g**. In all box plots, the center line represents median, the box edges show the 25^th^ and 75^th^ percentiles, and whiskers extend to maximum and minimum values.

To assess the degree of connectivity of GBM cells, we quantified the input neuron to starter GBM cell ratios, which were 20:1 for cortical transplantation sites and 2.3:1 for HIP transplantation (Fig. 2i). As a comparison, we derived SOX2^+^ neural progenitor cells (NPCs) from human iPSCs and transduced them with the RTG retrovirus (Extended Data Fig. 4f). Upon transplantation of ΔG rabies virus pre-infected NPCs into the RSP, we found a much lower input neuron to starter cell ratio (0.74:1 versus 18:1) and a lower number of total labeled neurons (2,300 versus 6,000) for NPCs compared to GBM cells transplanted in the same region at 10 dpt (Extended Data Fig. 4g-h). Estimated neuron to starter cell ratios from prior studies of transplantation of cortical organoids^23,38^ or NPCs^39–41^ into rodent brains were generally much lower than 1:1, indicating much higher neuronal connectivity of GBM cells in comparison to nonmalignant neural progenitor cells.

We also examined synaptic integration of GBM cells following longer term engraftment and extensive infiltration. We injected ΔG rabies virus one month after transplantation of GBM cells expressing RTG into the RSP (Extended Data Fig. 5a-b). At 10 days post rabies virus injection, starter DsRed^+^GFP^+^ GBM cells were distributed in the corpus callosum, RSP, and CA1 hippocampal regions, whereas neuronal inputs included both ipsilateral and contralateral cortices, ipsilateral thalamus, and basal forebrain (Extended Data Fig. 5b-c), with largely similar connection patterns to those found with pre-labeled GBOs (Fig. 2e, Extended Data Fig. 3b). Similarly, there was no evidence of glial labeling proximal to DsRed^+^GFP^+^ foci based on GFAP expression, further confirming the targeting specificity of this paradigm (Extended Data Fig. 5d). To rule out the possibility that cell death might drive leakage of G protein-expressing rabies virus to directly infect neurons, we stained for cleaved caspase 3 (cCas3) and did not observe significant apoptosis in DsRed^+^GFP^+^ GBM cells (Extended Data Fig. 5d).

Together, this systematic brain-wide connectome analysis revealed highly extensive and conserved integration into neuronal circuitry of diverse anatomical regions across the mouse brain with GBM cells from multiple patients, despite their inter- and intra-tumoral heterogeneity.

### Diverse neurotransmitter systems of synaptic inputs onto GBM cells

Next, we characterized the molecular identities of monosynaptic neuronal inputs onto GBM cells from diverse anatomical regions at 10 dpt. Simultaneous immunostaining for GFP^+^ rabies virus-labeled neurons and *in situ* hybridization for *vGLUT1/2* and *GAD1* revealed inputs from both glutamatergic and GABAergic neurons in the cortical, subcortical and hippocampal regions (Fig. 3a-c). We further found GFP^+^ cortical glutamatergic projections from both SATB2^+^ superficial layer neurons and CTIP2^+^ deep layer neurons (Extended Data Fig. 6a), while GFP^+^ GABAergic interneurons consisted of both PV^+^ and SST^+^ subtypes (Extended Data Fig. 6b). Quantification of cell type proportions across multiple transplantation sites revealed that glutamatergic (*vGLUT1/2*^+^*GAD1*^-^) inputs were by far the most abundant, and GABAergic (*vGLUT1/2*^-^*GAD1*^+^) and other (*vGLUT1/2*^-^*GAD1*^-^) subtypes exhibited area-specific differences in their connectivity (Fig. 3d).

**Figure 3.**
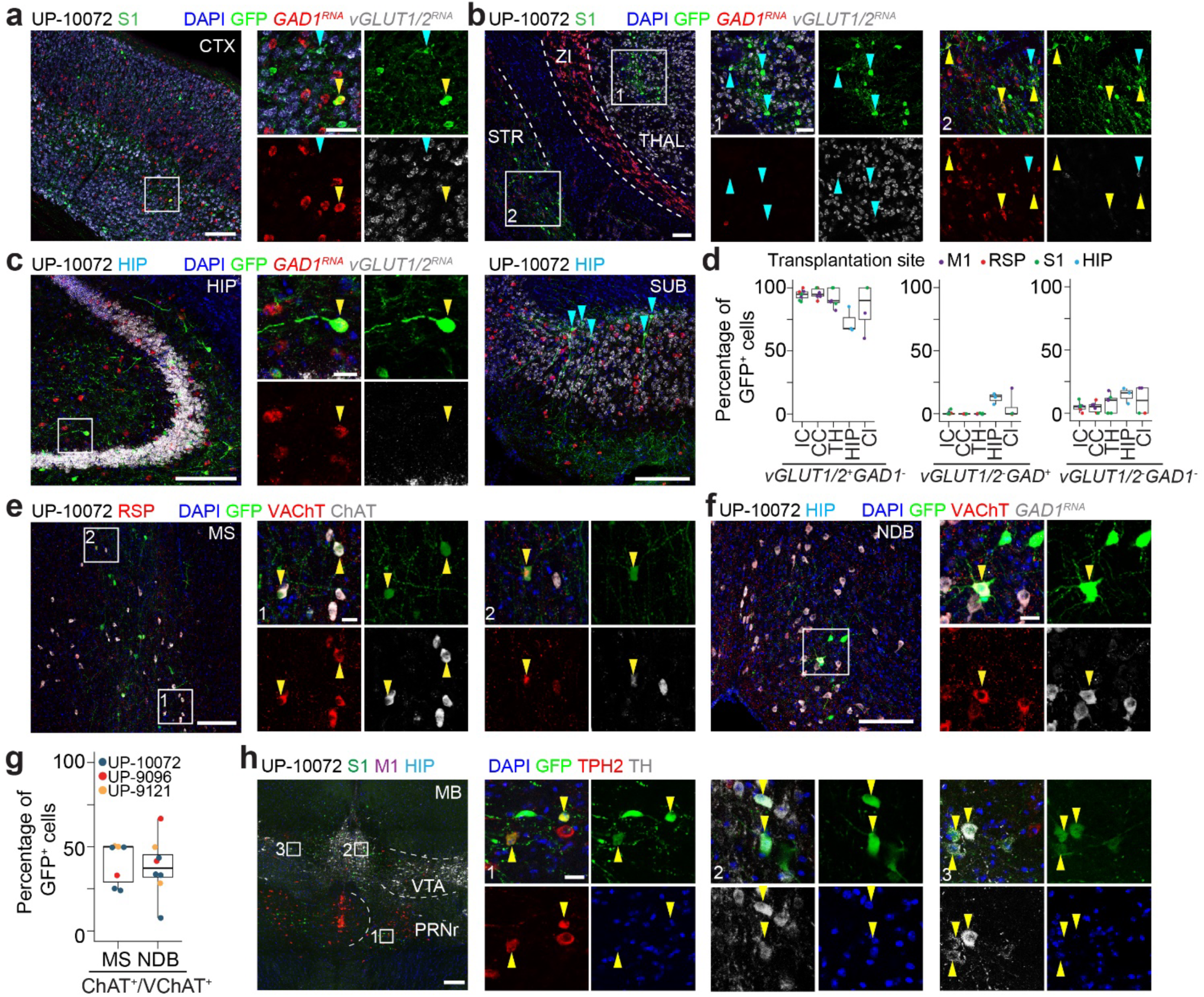
Integration of GBM cells into neuronal circuits with diverse neurotransmitter systems. **a-c**, Sample confocal images of RNA *in situ* hybridization for *GAD1* (red) and *vGLUT1/vGLUT2* (white) and GFP immunostaining of rabies virus-labeled neurons in the ipsilateral cortex (CTX) (**a**), ventral thalamus (inset 1) (**b**), striatum (inset 2) (**b**), and hippocampus (HIP)/subiculum (SUB) (**c**). Orange arrowheads indicate GABAergic neuron cell bodies, and blue arrows indicate glutamatergic neuron cell bodies. ZI, zona incerta. STR, striatum. **d**, Quantification of the percentages of GFP^+^ rabies virus-labeled neurons that were *vGLUT1/2*^+^*GAD1*^-^ (glutamatergic), *vGLUT1/2*^-^*GAD1*^+^ (GABAergic), or *vGLUT1/2*^-^*GAD1*^-^ (other) across brain regions. IC, ipsilateral cortex; CC, contralateral cortex; TH, thalamus; HP, hippocampus; Cl, claustrum. Dots are colored by transplantation site as indicated. Data are from *n* = 31 sections from *n* = 4 mice (from UP-10072 transplantations), with each dot representing one section. **e – f**, Sample confocal immunostaining images of VAChT^+^ChAT^+^GFP^+^ cholinergic neurons in the medial septum (MS, **e**) and diagonal band nucleus (NDB, **f**). Arrowheads indicate example cholinergic neurons. **g,** Quantification of the percentages of GFP^+^ neurons that were either VAChT^+^ or ChAT^+^ in MS or NDB. Data are from *n* = 7 mice (*n* = 4 RSP and *n =* 3 HIP transplantations) for MS and *n* = 8 mice (*n =* 2 HIP, *n* = 4 RSP, and *n* = 2 M1/S1/HIP transplantations) for NDB, with each dot representing one mouse, and color representing GBOs from different patients. In box plots in **d** and **g**, the center line represents median, edges represent 25^th^ and 75^th^ percentiles, and the whiskers extend to minimum and maximum values. **h**, Sample confocal images of TPH2^+^ serotonergic (inset 1) or TH^+^ dopaminergic (inset 2 and 3) GFP^+^ neurons in the midbrain (MB). VTA, ventral tegmental area; CS, superior central nucleus raphe; PRNr, pontine reticular nucleus. Arrowheads indicate example serotonergic or dopaminergic neurons of interest. For all sample images, GBO and transplantation location(s) are indicated with the same format as in Figure 2. Scale bars, 200 μm (low magnification images) and 20 μm (high magnification images).

We observed consistent projections from basal forebrain areas (MS and NDB) upon transplantation in RSP or HIP for GBOs across three patients (Fig. 2e). As many neurons from these basal forebrain areas are cholinergic neurons^42,43^, we performed immunostaining for ChAT and/or VAChT in MS (Fig. 3e) and NDB (Fig. 3f, Extended Data Fig. 6c, d), and found that about 40% of GFP^+^ neurons in these areas were indeed cholinergic (Fig. 3g). A subset of these GFP^+^ cholinergic neurons also expressed *GAD1*, denoting potential co-release of both acetylcholine (ACh) and GABA^44^ (Extended Data Fig. 6c, d). We additionally found sparse cholinergic inputs onto GBM cells from the brainstem pedunculopontine nucleus^45^ (Extended Data Fig. 6e). Several types of midbrain neuromodulatory neurons also projected onto GBM cells, including TPH2^+^ serotonergic neurons in the raphe nuclei^46^ and TH^+^ dopaminergic neurons in the ventral tegmental area^47^ (Fig. 3h, Extended Data Fig. 6f).

Together, our analyses revealed that, in addition to local and long-range glutamatergic and GABAergic synaptic inputs, there exist various long-range neuromodulatory projections of diverse neurotransmitter systems that may form synapses onto GBM cells. With the expression of receptors for these neurotransmitters in GBM (Fig. 1a), our results suggest extensive crosstalk between different neurotransmitter systems and GBM cells in the adult brain.

### Functional cholinergic synapses onto GBM cells mediated by metabotropic receptors

We next focused on cholinergic inputs from the basal forebrain onto GBM cells for more detailed analyses. Using expansion microscopy^48^, we confirmed localization of VAChT^+^ cholinergic presynaptic axon terminals adjacent to DsRed^+^ GBM cells in the RSP (Extended Data Fig. 7a). We also observed dense VAChT^+^ or ChAT^+^ puncta near EGFR^+^ tumor cells in primary IDH-wt GBM and IDH-mutant tumor tissue from several patients (Extended Data Fig. 7b-d).

To confirm a synaptic connection between cholinergic neurons and GBM cells using an independent transsynaptic viral tracing approach, we leveraged Cre-dependent and thymidine kinase (TK)-deficient herpes simplex virus (HSV, strain H129-LSL-ΔTK-tdTomato) for anterograde monosynaptic tracing in starter cells co-expressing Cre recombinase and TK^49,50^ (Fig. 4a). Co-injection of AAV-ChAT-Cre^51^ and AAV-DIO-TK-GFP into the basal forebrain resulted in expression of GFP in cholinergic axon terminals at distant locations, such as the RSP (Extended Data Fig. 7e). We next injected a mixture of H129-LSL-ΔTK-tdTomato, AAV-ChAT-Cre, and AAV-DIO-TK-GFP into the basal forebrain and simultaneously transplanted GBO cells in the HIP or RSP (Fig. 4a). By 6 dpt, we found HSV infection of GFP^+^ starter neurons with co-expression of GFP and tdTomato in the basal forebrain (Extended Data Fig. 7f). By 10 dpt, we found HSV infection of GBM cells with co-expression of tdTomato and human specific STEM121 or human nuclear antigen in the HIP (Fig. 4b, Extended Data Fig. 7g, h) or RSP (Extended Data Fig. 7i).

**Figure 4.**
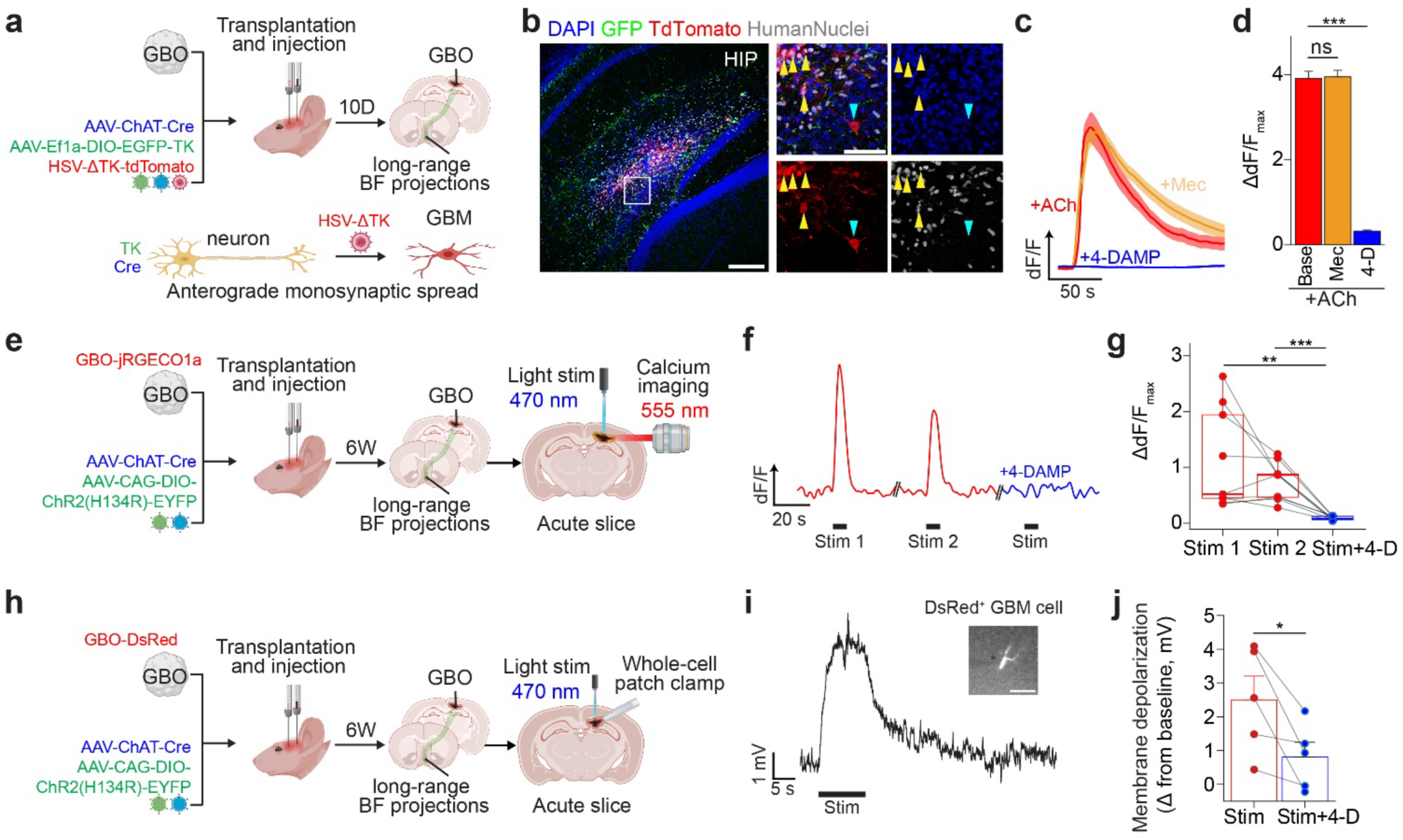
Direct functional cholinergic synapses onto GBM cells mediated by metabotropic receptors. **a**, Schematic illustrations of monosynaptic HSV anterograde tracing paradigm to confirm the projection from ChAT^+^ neurons in the basal forebrain onto GBM cells. BF: basal forebrain. **b**, Sample confocal images of TdTomato^+^HuNu^+^ (Human Nuclei) GBM cells (yellow arrowheads) in the hippocampus 10 dpt of GBO cells and viral injection to the basal forebrain. The blue arrowhead indicates an infected hippocampal neuron, and GFP^+^ fibers represent long-range basal forebrain projections. Images are representative of tracing experiments performed in *n* = 3 mice. Scale bars, 200 μm (low magnification images) and 20 μm (high magnification images). See additional images in **Extended Data** Fig. 7f-i. **c**, Representative traces of the Ca^2+^ response of UP-10072 GBOs to 1 mM acetylcholine (ACh) (red), 1 mM ACh in the presence of 100 μM 4-DAMP (blue), and 1 mM ACh in the presence of 100 μM mecamylamine (yellow). Data are plotted as mean ± s.e.m. of *n* = 10 cells from one representative organoid for each condition. **d**, Quantification of the maximal Ca^2+^ response to 1 mM ACh normalized to baseline intensity. Base, baseline; Mec, mecamylamine (100 μM); 4D, 4-DAMP (100 μM). Each bar represents data from *n* = 30 cells across *n =* 3 independent organoids. *P* = 0.9649 (baseline vs. Mec); *P* = 0.0001 (baseline vs. 4D); linear mixed-effects model fit by maximum likelihood with organoid number as a random variable; *p*-value adjustment for multiple comparisons with Tukey’s method. **e**, Schematic illustration of Ca^2+^ imaging paradigm of transplanted GBM cells in the acute slice. UP-10072 GBM cells expressing Ca^2+^ indicator JRGECO1a transplanted into the RSP with optogenetic stimulation of long-range basal forebrain projections (4-6 weeks post transplantation). **f**, Representative JRGECO1a fluorescence intensity trace of a GBM cell with three consecutive light stimulations following paradigm in **e**. Stimulation intervals are represented by black bars, and inter-recording intervals were at least 5 minutes. Red, stimulation in ACSF only; blue, stimulation in ACSF with 100 μM 4-DAMP. **g**, Quantification of maximal Ca^2+^ response of individual cells to light stimulation normalized to baseline intensity (*n* = 9 cells from 3 mice). Two-tailed paired Student’s *t*-test; FDR-adjusted *P* = 0.009 (Stim 1 vs. Stim + 4D), *P* = 7.3 x 10^-4^ (Stim 2 vs Stim + 4D). For the boxplot, center line represents median, edges represent 25^th^ and 75^th^ percentiles, and the whiskers extend to minimum and maximum values. **h**, Schematic illustration of electrophysiology experiments of transplanted GBM cells in the acute slice. UP-10072 GBM cells expressing DsRed transplanted into the RSP with light stimulation of long-range basal forebrain projections (4-6 weeks post transplantation). **i**, Representative GBM cell membrane depolarization in the current-clamp mode (I = 0 pA; resting membrane potential = −69 mV) in response to light stimulation in the presence of 25 μM CNQX and 200 μM 4-AP following paradigm in **h**. Patched DsRed^+^ GBM cell in the RSP (inset, right). Scale bar, 100 μm. **j**, Quantification of maximum membrane depolarization from baseline in response to light stimulation (*n* = 5 cells from 3 mice). Two-tailed paired Student’s *t*-test, *P* = 0.04. Bar plots in **d** and **j** are plotted as mean ± s.e.m. \**P* < 0.05, \*\**P* < 0.01, \*\*\**P* < 0.001.

To verify functional cholinergic synapses onto GBM cells, we performed calcium (Ca^2+^) imaging. We found that ACh exposure induced an immediate Ca^2+^ rise in GBOs derived from three patients, which was reduced by the M3 metabotropic receptor (CHRM3)-specific antagonist 4-DAMP, but not by pan-nicotinic antagonist mecamylamine (Fig. 4c-d, Extended Data Fig. 8a). To examine the response of GBM cells to synaptically-released ACh, we transplanted GBOs expressing red-shifted genetically-encoded Ca^2+^ indicator jRGECO1a^52^ in the RSP and simultaneously injected a combination of AAV-ChAT-Cre and AAV-DIO-ChR2(H134R)-EYFP into the basal forebrain (Fig. 4e). In acute slices from animals 6 weeks after transplantation, optogenetic stimulation of ChR2^+^ axon terminals induced Ca^2+^ transients in engrafted GBM cells, which were significantly attenuated by 4-DAMP (Fig. 4f, g, Extended Data Fig. 8b).

To further confirm direct functional cholinergic synaptic inputs onto GBM cells, we performed whole-cell patch-clamp electrophysiology. GBOs expressing DsRed were transplanted into the RSP and the combination of AAV-ChAT-Cre and AAV-DIO-ChR2(H134R)-EYFP was injected in the basal forebrain, and acute slices were prepared after 6 weeks (Fig. 4h). We confirmed inward currents in ChR2^+^ neurons following optogenetic stimulation (Extended Data Fig. 8c). Current-clamp recordings of GBM cells in the presence of AMPA receptor blocker CNQX revealed membrane depolarizations upon light stimulation, which were attenuated in amplitude by 4-DAMP (Fig. 4i, j, Extended Data Fig. 8d).

Together, our results define a functional long-range cholinergic synaptic input onto GBM cells mediated by the metabotropic CHRM3 receptor.

### Acute CHRM3 activation of GBM cells induces sustained calcium oscillations, transcriptional changes, and tumor invasion

To further examine CHRM3-dependent Ca^2+^ activity in response to ACh stimulation, we conducted Ca^2+^ imaging on GBOs in an air-liquid interface culture system^53^ with direct exposure to ACh. Consistent with prior reports^54^, we found that GBM cells in GBOs displayed periodic Ca^2+^ transients at a baseline mean frequency of 4.5 mHz (Fig. 5a). Brief (∼5 min) exposure of GBOs to ACh led to an increase in the frequency of spontaneous transients 30 minutes later with a mean frequency of 18 mHz (Fig. 5a-b, Extended Data Fig. 8e, f), which was significantly reduced by 4-DAMP (Extended Data Fig. 8g). These results demonstrate that ACh has not only immediate but also an extended influence on GBM cells, raising the possibility of additional functional effects such as modulation of distinct Ca^2+^ activity-dependent pathways^55^.

**Figure 5.**
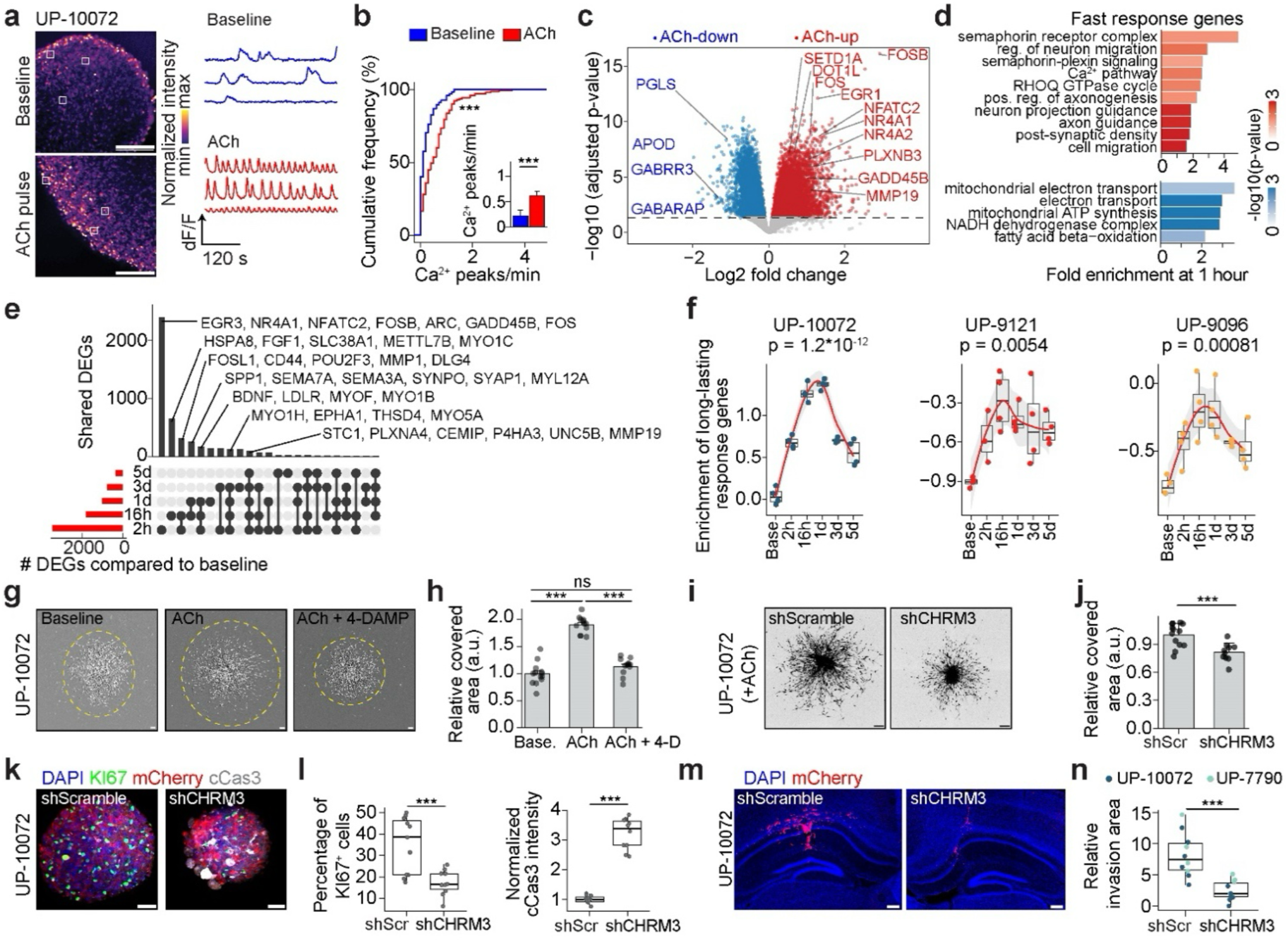
ACh-induced long-lasting Ca^2+^ oscillations, gene expression changes, and invasion of GBO cells via CHRM3. **a**, Representative Ca^2+^ imaging of UP-10072 GBOs under baseline conditions or 30 minutes after a pulse of 1 mM ACh on ALI. Insets (white squares) correspond to example cells in either condition with traces shown. Scale bars, 50 μm. **b**, Cumulative distribution plots of number of spontaneous Ca^2+^ peaks per minute in UP-10072 GBOs either under baseline conditions (blue) or 30 minutes after a 5-minute pulse of 1 mM ACh (red) (baseline, *n* = 122 cells from *n* = 3 GBOs; ACh, *n* = 140 cells from *n* = 3 GBOs). *P* = 2.8 x 10^-7^, Kolmogorov-Smirnov test. Inset, bar plot of Ca^2+^ peaks per minute, with data plotted as mean ± s.e.m. *P* = 1.0 x 10^-8^, linear mixed-effects model fit by maximum likelihood with GBO number as a random variable. **c**, Volcano plot of differentially expressed genes across UP-10072, UP-9096, and UP-9121 GBOs following 1-hour treatment of 1 mM ACh. Exemplary upregulated (red) and downregulated (blue) genes are indicated. Horizontal dashed line, adjusted *P*-value cutoff of 0.05 with effect size estimation by apeglm^96^. **d**, Representative GO terms for upregulated (red) or downregulated (blue) differentially expressed fast-response genes, with the x axis indicating fold enrichment of observed genes over expected. *P-*values, Fisher’s exact test, FDR *P* < 0.05. Reg., regulation; pos., positive. **e**, UpSet plot showing the co-occurrence of upregulated genes at various durations following a 1-hour pulse of ACh (2 hours, 16 hours, 1 day, 3 days, and 5 days). Exemplary genes for various DEG intersections are labeled. **f**, Time-dependent changes in the gene enrichment of long-lasting response genes to ACh. Each dot represents a distinct bulk transcriptomic sample. Curve represents the LOESS fit with the shaded area as s.e.m. *P* = 1.2 x 10^-12^ (UP-10072), *P* = 0.0054 (UP-9096), *P* = 0.00081 (UP-9121), one-way ANOVA. Base., baseline. **g – h**, Representative images (**g**) and quantification (**h**) of Matrigel matrix-based migration assay for UP-10072 GBOs, with dashed lines (yellow) representing the invaded area. Images were taken 48 hours following GBO seeding for baseline, 1 mM ACh, or 1 mM ACh with 100 μM 4-DAMP treatment. Scale bars, 200 μm. For quantification (**h**), the y-axis represents the mean covered area by GBO cells compared to baseline for n ≥ 12 GBOs per condition. One-way ANOVA with Tukey’s post hoc test, \**P* < 0.05, \*\*\**P* < 0.001. **i – j**, Representative confocal images (**i**) and migration area quantification (**j**) of UP-10072 GBOs with CHRM3 knockdown versus scrambled shRNA. Assays were performed in the presence of 1 mM ACh and images were taken 48 hours following GBO seeding (at least *n* = 10 organoids per condition). Data are plotted as mean ± s.e.m. Two-tailed Student’s *t-*test, *P* = 8.6 x 10^-4^. Scale bars, 200 μm. shScr, shScramble. **k**, Representative confocal immunostaining images of UP-10072 GBOs expressing either shScramble or shCHRM3 shRNAs 7 days after transduction, with mCherry representing shRNA expression. Scale bars, 50 μm. **l**, Quantification of the percentages of KI67^+^ cells and normalized cleaved caspase 3 (cCas3) intensity in shScramble (*n* = 11) vs shCHRM3 (*n* = 10) GBOs. Two-tailed Student’s *t*-test, *P* = 9.9 x 10^-4^ (KI67 percentage), *P* = 2.8 x 10^-11^ (cCas3 intensity). Norm., normalized. **m**, Representative confocal images of UP-10072 GBOs expressing either shScramble or shCHRM3 3 weeks post transplantation into the hippocampus, with mCherry representing shRNA expression. Scale bars, 200 μm. **n**, Quantification of relative invasion area of shScramble or shCHRM3 GBM cells 3 weeks post transplantation (UP-10072, *n* = 6 mice, HIP; UP-7790, *n* = 2 mice, striatum; UP-7790, *n* = 2 mice, HIP). Two-tailed Student’s *t-*test, *P* = 1.8 x 10^-4^. For all box plots, the center line represents median, edges represent 25^th^ and 75^th^ percentiles, and the whiskers extend to minimum and maximum values. \*\*\**P* < 0.001.

We next examined ACh-induced transcriptional changes in GBM cells with bulk and single-cell RNA sequencing analyses. We devised a time-resolved bulk RNA sequencing paradigm to explore transcriptional dynamics upon continuous ACh stimulation or in response to varying pulse durations of ACh with sequencing at 1 hour (Extended Data Fig. 9a). Analysis of GBOs from three patients revealed ACh-induced upregulation of immediate early genes (such as AP-1 family transcription factors *FOS* and *FOSB*, *EGR1*), nuclear transcription factors associated with Ca^2+^ oscillations (such as *NFATC2* at ∼20 mHz^55^), epigenetic regulators (such as *GADD45B, DOT1L,* and *SETD1A*), and genes related to axon guidance and motility (such as *PLXNB3* and *MMP19*) (Fig. 5c). Many of these ACh-induced genes exhibited a time-dependent increase in expression levels over the 1-hour period (Extended Data Fig. 9b), whereas brief (5, 15, or 30 minute) pulses of ACh followed by washout and analysis at 1 hour revealed equivalent levels of upregulation (Extended Data Fig. 9c), suggesting rapid induction of transcriptional reprogramming driven by acute ACh exposure. We defined genes with upregulated expression at 1 hour as fast-response genes, which were enriched in Gene Ontology (GO) terms related to semaphorin-plexin signaling, cellular migration, and post-synaptic density, while downregulated genes were generally associated with metabolic function (Fig. 5d). Gene signature enrichment showed significant upregulation of the AP-1 FOS transcription factor family, migration, and axon guidance gene sets across GBOs from different patients at 1 hour (Extended Data Fig. 9d-f). We also conducted scRNAseq analyses of these GBOs after 1-hour treatment with ACh, which confirmed upregulation of the fast-response gene signature (Extended Data Fig. 9g-j). At the single-cell level, GBM cells more highly enriched in post-synaptic density genes tended to also be more enriched in ACh fast-response genes (Extended Data Fig. 9k). In addition, we found that higher expression of this fast response signature was correlated with worse patient prognosis in GBM cohorts from The Cancer Genome Atlas^56^ (TCGA) and the Chinese Glioma Genome Atlas^57^ (CGGA) (Extended Data Fig. 9l).

We then asked whether brief exposure of GBOs to ACh would be sufficient to exert long-lasting transcriptional changes beyond 1 hour. We performed bulk RNA sequencing after various lengths of time following a 1-hour pulse of ACh (Extended Data Fig. 10a). At day 1, we found significant changes in the expression of many genes, which we defined as long-lasting response genes (Extended Data Fig. 10b). These genes were enriched in GO terms related to cellular adhesion, contractility, and migration (Extended Data Fig. 10c). Enrichment of the long-lasting response gene signature gradually decreased over time, but remained elevated at day 5 (Fig. 5e, f). Genes such as *STC1, PLXNA4, MMP19, UNC5B, CEMIP*, and *P4HA3*, which are known to play roles in invasion^58,59^ and progression^60^ of glioma or other tumors^61–63^, remained upregulated for 5 days following 1-hour ACh exposure (Fig. 5e). Accordingly, enrichment of this long-lasting response signature was also associated with decreased patient survival time reported in the TCGA and CGGA datasets (Extended Data Fig. 10d).

To confirm the functional impact of ACh-induced transcriptional changes associated with cellular phenotypes such as migration, we performed invasion and migration assays. We found that in an all-human cell assembloid model, a single 1-hour pulse of ACh pretreatment was sufficient to increase GBO cell invasion into human iPSC-derived sliced neocortical organoids^28^ over 2 days (Extended Data Fig. 11a-b). We also applied a Matrigel matrix-based assay to examine GBM cell migration across GBOs derived from 5 different patients in response to ACh treatment and subsequent inhibition of CHRM3 (Fig. 5g-h, Extended Data Fig. 11c-d). While we found patient-specific variability in the degree to which ACh increased migration in this assay, 4-DAMP uniformly reduced migration compared to ACh stimulation in GBOs from all patients (Fig. 5g-h, Extended Data Fig. 11c-d). The heterogeneity in ACh-induced migration combined with consistent inhibition by 4-DAMP, even to levels below baseline in some patients, may be explained by cell-intrinsic constitutive activity of the CHRM3 G-protein coupled receptor^64,65^ or activation of this receptor by other metabolites, such as choline present in the culture medium^66,67^.

To explore CHRM3 as a molecular target for GBM, we expressed short-hairpin RNA (shRNA) targeting CHRM3 in GBOs via lentivirus (Extended Data Fig. 11e). CHRM3 knockdown in GBOs decreased the transcriptional response to ACh as assayed by RNA sequencing (Extended Data Fig. 11f). Knockdown of CHRM3 significantly inhibited GBO migration *in vitro* in the presence of ACh compared to a scrambled control shRNA after 48 hours (Fig. 5i-j, Extended Data Fig. 11g-h). After 7 days post transduction, we observed a significant decrease in the GBO size and percentage of KI67^+^ cells as well as an increase in cCas3 expression, indicating a negative impact on cell proliferation and survival (Fig. 5k-l, Extended Data Fig. 11i-j). CHRM3 knockdown GBOs also exhibited decreased tumor cell invasion into neocortical organoids in the all-human cell assembloid model with 1-hour ACh pre-treatment (Extended Data Fig. 11k-l). We further transplanted CHRM3 knockdown GBOs into adult immunodeficient mice and found a significant decrease in tumor burden and invasion area compared to GBOs expressing a scrambled control shRNA (Fig. 5m-n).

Taken together, these results indicate that acute ACh stimulation of GBM cells can lead to a sustained impact on GBM cells through CHRM3 and that blockade of CHRM3 function leads to decreased survival, proliferation, and invasion of GBM cells.

## DISCUSSION

By applying monosynaptic viral tracing to systematically define malignant neuronal circuits, our study demonstrates rapid establishment of brain-wide connectivity of GBM cells into diverse neurotransmitter networks. Beyond the pioneering discoveries of local glutamatergic inputs onto glioma cells that signal through ionotropic receptors^7,8,15^, our study reveals strikingly extensive interactions between GBM cells and neurons that release different neurotransmitters and originate in different brain regions, including both bilateral cortices and subcortical areas, such as the basal forebrain, brainstem and thalamic nuclei. Given that rabies virus does not label all synaptic inputs of starter cells^68,69^, our observation may still represent an underestimation. Our findings thus significantly expand the notion that tumor cells may be innervated and regulated by myriad neuronal subtypes^70–73^, such as those comprising neuromodulatory systems^74,75^ that signal through metabotropic neurotransmitter receptors. Given the ability of glioma to bidirectionally interact with neuronal circuits^19,20,76,77^ combined with established roles of diffuse neuromodulatory projections in memory and behavior^43,78^, our discoveries also provide a foundation to investigate whether feedback to neurons that innervate GBM cells might explain generalized patient symptoms such as cognitive dysfunction, sleep disturbances, seizures, or behavioral deficits that affect quality of life^79–82^ and to develop corresponding therapeutic interventions.

Employing cholinergic inputs from the basal forebrain as an example, we validated functional cholinergic synapses onto GBM mediated by the M3 metabotropic receptor. Downregulation of CHRM3 showed the potential to attenuate GBM progression *in vitro* and *in vivo*. Importantly, we discovered a long-lasting impact of brief ACh stimulation on GBM cells, defined by transcriptional reprogramming and enhanced migratory behavior. Given that GBM cells are highly invasive, our results collectively support a model in which rapid formation and acute activation of synapses onto migratory GBM cells may refuel tumor cells to promote their migration, survival, and progression, analogous to gas stations refueling cars traveling along the highway.

Together, our study reveals diverse and robust neuronal inputs onto GBM cells, and our findings may serve as a framework to investigate the functional impact and therapeutic relevance of distinct synaptic inputs onto glioma.

## METHODS

### Human specimens and animal models

De-identified human GBM surgical samples were collected at the Hospital of the University of Pennsylvania after informed patient consent under a protocol approved by the Institutional Review Board of the University of Pennsylvania. Sample distribution and collection were overseen by the University of Pennsylvania Tumor Tissue/Biospecimen Bank in accordance with ethical and technical guidelines on the use of human samples for biomedical research. Primary and recurrent GBM specimens were included in this study. Epidemiological data for each subject and genomic data were provided by the Neurosurgery Clinical Research Division (NCRD) at the University of Pennsylvania. Disease-relevant genomic alterations (Agilent Haploplex assay, Illumina HiSeq2500), fusion transcripts (Illumina HiSeq2500), and MGMT promoter methylation (PyroMark Q24, Qiagen) were performed by the University of Pennsylvania Center for Personalized Diagnostics.

All animal experiments in this study were conducted in accordance with institutional guidelines and protocols approved by the Institutional Animal Care and Use Committee (IACUC) at the University of Pennsylvania. Animals were housed on a 12-hour light/dark cycle with food and water *ad libitum*. We used 5–8-week-old female athymic nude mice (*Foxn1^nu^/Foxn1^nu^*, Jackson Laboratory, Strain #007850) for all experiments. Animals were monitored routinely for weight loss and physical/neurological abnormalities.

### GBM organoid culture

GBM cells used were cultured as 3D GBM organoids (GBOs), which were generated directly from human GBM surgical specimens following our established protocols^25,83^. In brief, fresh surgically resected GBM tissue was placed in dissection medium consisting of Hibernate A (Thermo Fisher, A1247501), 1X GlutaMax (Thermo Fisher Scientific, 35050061), and 1X Antibiotic-Antimycotic (Thermo Fisher Scientific, 15240062) and kept at 4°C. Tissue was transported to the lab and subsequently dissected into ∼1 mm^3^ pieces with small spring scissors in a sterile petri dish. Dissected tumor pieces were washed in 1X Red Blood Cell (RBC) Lysis Buffer (Thermo Fisher Scientific, 00-4222-57) and then washed with DPBS (Thermo Fisher Scientific, 14040182). Samples were then transferred to an ultra-low attachment 6-well culture plate to be cultured in GBO culture medium containing 50% Neurobasal (Thermo Fisher Scientific, 21103049), 50% DMEM:F12 (Thermo Fisher Scientific, 11320033), 1X NEAAs (Thermo Fisher Scientific, 11140050), 1X GlutaMax (Thermo Fisher Scientific, 35050061), 1X Penicillin-Streptomycin (Thermo Fisher Scientific, 15070063), 1X B27 without vitamin A supplement (Thermo Fisher Scientific, 12587010), 1X N2 supplement (Thermo Fisher Scientific, 17502048), 1X 2-mercaptoethanol (Thermo Fisher Scientific, 21985023), and 2.5 μg/ml human recombinant insulin (Sigma, 19278). Wide-bore P1000 pipette tips with ∼3 mm diameter openings were used to transfer tumor pieces. 6-well plates containing GBOs were cultured on an orbital shaker with continuous shaking speed (110 rpm) in a 37°C, 5% CO_2_, and 85% humidity sterile culture incubator, and culture medium was replaced every 2 days. Generally, during the first 1-2 weeks of culture, cellular debris shed from tumor pieces could be observed; however, round GBOs typically formed after 2-3 weeks. GBOs were passaged by cutting larger (>1 mm^3^) pieces to approximately 0.5 – 1 mm^3^ pieces with dissection scissors. Biobanking and long-term storage of GBOs^25,83^ was performed by transferring small GBO pieces (0.1 – 0.2 mm^3^) to GBO culture medium containing 10% DMSO in cryogenic vials (Thermo Fisher Scientific, 13-700-504). Vials were stored in a foam Cell Freezing Container (Thermo Fisher Scientific, 07-210-002) at −80°C overnight and then transferred to liquid nitrogen for long-term storage.

### Human iPSC-derived progenitor cell and organoid culture

Human neural progenitor cells were derived from the dissociation of human iPSC-derived forebrain organoids generated following a protocol reported previously with minor modifications^84^. In brief, detached WTC-11^85^ human iPSC cells were transferred to an ultra-low attachment U-bottom 96-well plate (20K cells/well) and cultured in mTeSR Plus media (StemCell Technologies, 5825) supplemented with 10 μM Y-27632 (StemCell Technologies, 72304) for 48 hours to achieve Embryoid Body (EB) aggregation. On days 3-7, EBs were cultured in F1 neural induction medium containing DMEM/F12 supplemented with 20% KnockOut Serum Replacement (Thermo Fisher Scientific, 10828028), 1X Penicillin-Streptomycin, 1X NEAAs, 1X GlutaMax, 0.1 mM 2-mercaptoethanol, 0.0002% heparin, 1 μM IWR-1-endo (StemCell Technologies, 72562), 5 μM SB-431542 (StemCell Technologies, 72234), and 1 μM LDN-193189 (StemCell Technologies, 72147). On day 7, organoids were embedded in Matrigel (Corning, 8774552) and cultured in F2 medium containing DMEM/F12 supplemented with 1X N2 supplement, 1X Penicillin-Streptomycin, 1X NEAAs, 1X GlutaMax, 0.1 mM 2-mercaptoethanol, 1 μM SB-431542, and 1 μM CHIR99021 (StemCell Technologies, 72054) for 7 days. On day 14, embedded organoids were dissociated from Matrigel and transferred to an ultra-low attachment 6-well plate, placed on an orbital shaker at 110 rpm, and cultured in F3 medium containing 50% DMEM/F12 and 50% Neurobasal supplemented with 1X B27 supplement (Thermo Fisher Scientific, 17504044), 1X N2 supplement, 1X Penicillin-Streptomycin, 1X NEAAs, 1X GlutaMax, 0.1 mM 2-mercaptoethanol, and 3 μg/ml human insulin. On day 20, the forebrain organoids were digested with Accutase (Thermo Fisher Scientific, A1110501) at 37°C for 15 minutes and dissociated to single cells, which were later seeded on plates pre-coated with 1% Matrigel and cultured in F3 medium supplemented with 1 μM CHIR-99021, bFGF (20 ng/mL, PeproTech, 100-18B), and EGF (20 ng/mL, PeproTech, AF-100-15).

Human iPSC-derived sliced neocortical organoids (SNOs) were generated and maintained as described above for neural progenitor cells using either the WTC-11^85^ or C1-2^84^ iPSC lines but with the following modifications: organoids were not dissociated at day 20 but rather maintained in culture until day 70, at which point organoids were sliced as previously described^28^ and cultured in F4 medium containing Neurobasal supplemented with 1X B27 supplement, 1X GlutaMax, 1X NEAAs, 1X 2-mercaptoethanol, 1X Penicillin-Streptomycin, 0.05 mM cAMP (STEMCELL Technologies, 73886), 0.2 mM ascorbic acid (Sigma, A0278), 20 ng/mL BDNF (PeproTech, 450-02), and 20 ng/mL GDNF (PeproTech, 450-10) thereafter.

### Single-cell and bulk RNA sequencing

For single-cell RNA sequencing (scRNAseq) library preparation, we employed a plate-based scRNAseq method based on SMART-seq3^27,86^ with minor modifications. GBOs were first dissociated into a single-cell suspension with a brain tumor dissociation kit (Miltenyi Biotech, 130-0950929). In brief, 5-10 GBOs were washed with DPBS and incubated in 1 mL of dissociation mix according to the manufacturer’s protocol. GBOs were placed on a tube rotator in a 37°C incubator for 30 minutes to 1 hour with occasional pipetting to mechanically disrupt large tissue chunks. Cells were then strained through a 70 μm filter (Miltenyi Biotech, 130-110-916), centrifuged at 300g for 5 minutes and resuspended in 1 mL of GBO culture medium. Cell viability and cell concentration were measured with an automated cell counter (Thermo Fisher Scientific, Countess 3 Automated Cell Counter) with trypan blue staining (Thermo Fisher Scientific, C10312). An ideal cell viability following dissociation is typically > 80%. Cells were then resuspended in FACS pre-sort buffer (BD, 563503) with 0.2 μg/mL DAPI (BD, 564907). For sliced neocortical organoid (SNO) scRNAseq, 100-day old SNOs cultured as described above were dissociated in a similar manner as GBOs but with the addition of 50 μL Enzyme P (Miltenyi, 130-107-677) per 1 mL of dissociation mix. Single cells were sorted on a BD Influx (100 μm nozzle) with FACSDiva software into low-profile 96-well PCR plates (USA Scientific, 1402-9500) containing 3 μL of SMART-seq3 lysis buffer (0.5 μL PEG 8000 (40% solution, Sigma Aldrich, P1458), 0.03 μL Triton X-100 (10% solution), 0.02 μL of 100 μM Oligo-dT30VN, 0.2 μL of 10 mM dNTPs (Roche, 50-196-5273), 0.02 μL Protector RNase inhibitor (Sigma Aldrich, 3335402001), 0.02 μL recombinant RNase inhibitor (Takara, 2313B) and 2.21 μL nuclease-free water per reaction (Thermo Fisher Scientific, AM9932)) and 3 μL Vapor-Lock (Qiagen, 981611). Plates after sorting were briefly centrifuged, frozen on dry ice, and stored at - 80°C for later processing.

For library preparation, plates were thawed, heated to 72°C for 10 minutes, and subsequently kept at 4°C for cell lysis. 3 μL of RT mix, containing 0.1 μL of 1 M Tris-HCl pH 8.3 (Hampton Research, HR2-900-14), 0.024 μL of 5 M NaCl (Thermo Fisher Scientific, AM9759), 0.01 μL of 1 M MgCl_2_ (Thermo Fisher Scientific, AM9530G), 0.04 μL of 1 mM GTP (Thermo Fisher Scientific, R1461), 0.32 μL of 100 mM DTT (Millipore Sigma, 3483-12-3), 0.016 μL of nuclease-free water, 0.025 μL Protector RNase inhibitor (Millipore Sigma, C852A14), 0.025 μL recombinant RNase inhibitor (Takara, 2313B), 0.04 μL reverse transcriptase (Thermo Fisher Scientific, EP0753), and 0.4 μL of 20 μM TSO, was added to each well of the 96-well plate. To enable multiplexing and enhance throughput, we designed 48 unique TSOs consisting of an additional 6-bp cell barcode directly adjacent to the 5’-end of the Smart-seq3 UMI. Reverse transcription was performed with the following thermocycling conditions: 42°C for 90 minutes, 10 cycles of 50°C for 2 minutes and 42°C for 2 minutes, 85°C for 5 minutes, and 4°C indefinitely. Next, for cDNA amplification, 6 μL of PCR mix, consisting of 2 μL 5X KAPA HiFi HotStart Buffer (Roche, KK2502), 0.048 μL of 10 mM dNTPs, 0.005 μL of 1 M MgCl_2_, 0.1 μL of 100 μM PCR forward primer, 0.01 μL of 100 μM PCR reverse primer, 3.64 μL of nuclease-free water, and 0.2 μL of KAPA HiFi HotStart polymerase (Roche, KK2502), was added to each well. cDNA amplification was performed with the following thermocycling conditions: 98°C for 3 minutes, 18 cycles of 98°C for 20 seconds then 65°C for 30 seconds then 72°C for 4 minutes, 72°C for 5 minutes, and 4°C indefinitely. cDNA from 48 wells was then pooled together into one sample (2 samples per plate), and samples were then purified twice with 0.6X then 0.8X AMPure XP beads (Thermo Fisher Scientific, A63881) and eluted in 10 μL of nuclease-free water. cDNA was quantified using Qubit dsDNA HS assay kit (Life Technologies, Q32851).

Each sample was then tagmented by mixing 2 μL of 0.5 ng/μL cDNA, 4 μL of 2X TD buffer (20 mM Tris pH 8.0, 10 mM MgCl_2_, 20% dimethylformamide (Sigma, D4551)), and 2 μL of Tn5 (Thermo Fisher Scientific, TNP92110) and incubating at 55°C for 20 minutes with reaction termination upon addition of 2 μL of 0.2% SDS (Invitrogen, 15553-035) at room temperature for 5 minutes. We chose to use i5-only Tn5 in order to enrich for the 5’-ends of mRNA, although full-length sequencing can be achieved using i5/i7 Tn5. Fragments were amplified by adding 1 μL of 10 μM Nextera i7 primer, 1 μL 10 μM Nextera i5 primer, 5 μL of 5X Phusion Plus Buffer (Thermo Fisher Scientific, F630L), 0.5 μL of 10 mM dNTPs, 0.5 μL of 1% Tween-20, 6.8 μL nuclease-free water, and 0.2 μL of Phusion Plus polymerase (Thermo Fisher Scientific, F630L). Tagmentation PCR was performed with the following thermocycling conditions: 98°C for 3 minutes, 8 cycles of 98°C for 10 seconds then 60°C for 30 seconds then 72°C for seconds, 72°C for 5 minutes, and then indefinitely at 4°C. Resulting DNA was purified twice with 0.8X then 1.0X AMPure XP beads and eluted in 10 μL of 10 mM Tris, pH 8.0. Samples were then quantified by qPCR with library sizes quantified by Bioanalyzer (Agilent). Samples were sequenced on either a NovaSeq 6000 (Illumina) or NextSeq 550 (Illumina) at a final depth of roughly 100 K-200 K reads/cell.

For bulk RNA sequencing, GBOs were reaggregated into organoids of 2000 cells in 96-well U-bottomed plates as described above. For ACh experiments, ACh (acetylcholine chloride, Sigma Aldrich, A6625) was applied to a final concentration of 1 mM for various durations prior to removal of medium and sample lysis. If any condition required a waiting period prior to lysis, GBOs were washed once with GBO medium prior to complete replacement with fresh GBO medium. For sequencing of CHRM3 knockdown GBOs, GBO cells were directly infected with either scrambled or knockdown lentivirus prior to reaggregation as described above for retroviral infection, and cells were subjected to 1-hour ACh treatment 7 days following shRNA transduction. Library preparation was performed as described above for single-cell sequencing with minor modifications: all reactions were scaled up by 10X the volume (e.g., sample lysis in 30 μL of Smart-seq3 lysis buffer), cDNA amplification was performed with 16 cycles, and tagmentation PCR was performed with 10 cycles. Libraries were sequenced to roughly 12-15 M reads per sample.

### Sequencing pre-processing and analysis

To process single-cell RNA sequencing data, raw files were demultiplexed with bcl2fastq (Illumina) without adapter trimming and with the option for *--create-fastq-for-index-reads*. A combined fastq file consisting of a 22-bp cell barcode (composed of 8-bp index 1, 8-bp index 2, and 6-bp barcode) was then generated. Alignment was performed using STARsolo as part of STAR v2.7.10b^87,88^ with GRCh38 as the reference genome and gencode v.41 GTF as the annotation file, and with the following additional parameters: *--alignIntronMax 1000000 --outFilterScoreMinOverLread 0.3 --outFilterMatchNminOverLread 0.3 --limitOutSJcollapsed 4000000 --soloType CB_UMI_Simple --soloCBstart 1 --soloCBlen 22 --soloUMIstart 23 --soloUMIlen 8 --soloBarcodeMate 1 --clip5pNbases 30 --soloCBmatchWLtype 1MM_multi --soloCellFilter EmptyDrops_CR --soloStrand Reverse*. A cell barcode text file was supplied, and multimapping alignments were discarded.

Count matrices via the “GeneFull” option including intronic counts from STARsolo were imported into R (v4.3.1) using Seurat (v4.3.0.1)^89^. Genes expressed in less than 10 cells were discarded, and cells that had less than 1000 UMIs or with a percentage of mitochondrial UMIs over 20% were discarded as well. Counts were normalized with SCTransform^90^ with vst.flavor = “v2”, variable.features.n = 15000, and regression of percentage mitochondrial UMIs and number of UMIs. GBM cellular state was assigned as previously described in^13^ and implemented in^91^ in the function get.sig.scores. Relative enrichment scores of various gene signatures were computed via UCell^92^ package as implemented in the function AddModuleScore_UCell. To identify clusters in the SNO dataset, the FindAllMarkers function in Seurat was used with adjusted *p*-value < 0.05 (Wilcoxon rank-sum test with Bonferroni correction) and log fold-change threshold of 0.25. We also retrieved count matrices for various published datasets using either the Smart-seq2 platform^13,30^ or 10X Genomics platform^29^. Annotated tumor cells from 6 patients (SF10108, SF11082, SF11780, SF12382, SF3391, SF9510) were merged from the Wang et al.^29^ dataset. The Neftel et al.^13^ dataset was first normalized by SCTransform^90^ and then integrated using Harmony^93^. Dot plots in Figure 1a and Extended Data Figure 1j were plotted with the ‘RNA’ assay and ‘data’ slot following log-normalization with the dittoDotPlot function implemented in the dittoSeq package^94^. scRNAseq data of GBOs at baseline conditions or treated with ACh were first normalized with SCTransform^90^ with method = ‘glmGamPoi’ and then integrated with rPCA in Seurat^89^.

Bulk RNA sequencing data with UMIs were processed similarly but with an additional option of *--soloUMIdedup Exact* during alignment to account for increased UMI complexity of these samples. Samples with at least 10000 detected genes were included for further analysis. For the 1 hour ACh pulse experiments (fast response, Extended Data Fig. 9a-f, Fig. 5c-d), differential expression analysis was performed with DESeq2^95^ using UMI counts as input and a model design of *∼Tumor + Treatment* (where *Treatment* indicates whether ACh was applied and *Tumor* indicates patient GBO) to control for differences amongst patients, with effects size estimation performed with apeglm^96^. For the pulse experiments for long-lasting transcriptional changes (long-lasting response, Extended Data Fig. 10, Fig. 5e-f), differential expression analysis was performed as above but with a model design of ∼*Tumor + Condition* (where *Condition* represents the duration prior to sample lysis). Differentially expressed genes between the control (no ACh) condition and each of the durations were obtained separately (adjusted *p*-value < 0.05) and visualized with the UpSetR package^97^. Gene Ontology (GO) analyses of differentially expressed genes were performed with an overrepresentation test as implemented in pantherdb.org^98^ with sets of either upregulated or downregulated genes (defined as adjusted *p*-value < 0.05 and abs(log_2_FC) > 0.25). The background genes for GO analysis were defined as genes that were detected with a low threshold (average of 0.25 – 0.5 UMIs per sample). The ACh fast response gene signature was defined as the top 100 genes upregulated from baseline after 1-hour ACh treatment ranked by adjusted *p*-value. The ACh long-lasting response gene signature was defined as the top 100 genes upregulated from baseline 1 day after 1-hour pulse of ACh ranked by adjusted *p*-value. Exemplary genes in Extended Data Fig. 9b-c were plotted as a fold-change from baseline using the DESeq2-normalized UMIs. Gene enrichment for various pathways was obtained by importing the UMI count matrix into Seurat (with each ‘cell’ as a bulk sample), running SCTransform with vst.flavor = “v2”, and using the AddModuleScore function with the ‘SCT’ assay as implemented in Seurat.

### Viral vector and plasmid generation

EnvA-pseudotyped G-deleted EGFP rabies virus was purchased from the Salk viral core (32635). Monosynaptic HSV (strain H129-LSL-ΔTK-tdTomato) was a kind gift from Lynn Enquist and expanded in-house. AAV2/9-ChAT-Cre-WPRE-hGHpA (PT-0607) and AAV2/8-EF1α-DIO-EGFP-2a-TK-WPRE-pA (PT-0087) were purchased from BrainVTA. AAV9-EF1α-DIO-hChR2(H134R)-EYFP-WPRE-hGHpA was a gift from Karl Deisseroth (Addgene, viral prep 20298-AAV9). The retroviral CAG-dsRed-T2A-RabiesG-IRES-TVA (RTG helper plasmid) was a kind gift from Benedikt Berninger. To generate the control retroviral helper plasmid without G protein, we excised the G protein coding sequence and ligated the plasmid with an annealed duplex oligonucleotide. Plasmid sequences were confirmed by Sanger DNA sequencing (Penn Genomic and Sequencing Core) and whole plasmid sequencing (Plasmidsaurus).

ShRNA sequences for CHRM3 were designed using Broad Institute RNAi Consortium (https://www.broadinstitute.org/rnai-consortium/rnai-consortium-shrna-library). The backbone vector for shRNAs was purchased from Addgene (pLKO.1_mCherry, 128073). We followed the shRNA construction protocol from the Genetic Perturbation Platform web portal (https://portals.broadinstitute.org/gpp/public/resources/protocols). Plasmid sequences were confirmed by Sanger DNA sequencing.

Retroviruses were produced with HEK 293T cells. 293T packaging cells were prepared at 70-80% confluency in 15 cm tissue culture plates. For each plate, 6 μg pMD2.G (Addgene, 12259), 4 μg pUMVC (Addgene, 8449) and 18 μg transfer plasmid were mixed in 700 μL DMEM medium (Corning, 10-013-CV). Subsequently, 84 μL of LipoD293^TM^ (SignaGen Laboratories, SL100668) was mixed with 700 μL DMEM medium. These two mixtures were then combined and incubated at room temperature for 10 minutes. The plate was then replenished with 15 mL of pre-warmed 293T culture medium (DMEM, 10% fetal bovine serum (Fisher Scientific, SH3007103) and 1X Penicillin-Streptomycin) and the combined mixture was added dropwise. Virus-containing medium was collected after 24, 48 and 72 hours and stored at 4°C. The 293T culture was replenished with 15 mL warm culture medium after each collection. After 72 hours, the virus-containing medium was pooled together and centrifuged at 500g for 5 minutes to pellet cells. The supernatant was filtered through a 0.45 μm PES filter (Thermo Fisher Scientific, 566-0020). The filtered medium was then centrifuged with a high-speed centrifuge at 25,000g and 4°C for 2 hours followed by resuspension in DPBS. Concentrated viruses were aliquoted and stored at −80°C. Viral titer was determined by serial dilution, infection in 293T cells, and counting of positive colonies. Lentiviruses were produced using a similar procedure with 293T cells using the psPAX2 plasmid (Addgene, 12260) instead of pUMVC.

### Stereotaxic GBO transplantation and virus injection

For all transplantation experiments, GBO cells were prepared in a single-cell suspension via the dissociation protocol as described above and kept in sterile Hibernate A prior to the surgical procedure. Transplantation was performed aseptically following IACUC guidelines for rodent survival surgery. The cranium was exposed following a midline scalp incision, and a hole was drilled through the cranium above the desired stereotaxic coordinates using a micromotor drill (Stoelting, 51449). Roughly 200,000 single GBO cells in 1-2 μL of Hibernate A were loaded into a 5 μL Hamilton syringe (Hamilton, 80016) with a 26-gauge needle (Hamilton, 7768-02). Injection stereotaxic coordinates used for monosynaptic tracing were as follows: M1 (1.2 mm anterior to bregma (A/P), 1.5 mm lateral to midline (M/L), and 1.5 mm deep to cranial surface (D/V)), S1 (A/P - 0.9 mm, M/L +3.0 mm, D/V −1.5 mm), RSP (A/P −2.3 mm, M/L 0.4 mm, D/V −1.7 mm), ventral hippocampus (A/P −2.0 mm, M/L +1.6 mm, D/V −2.6 mm), and dorsal hippocampus (A/P −2.0 mm, M/L +1.6 mm, D/V −1.8 mm). Injection was performed with a flow rate of less than 0.2 μL/min. The needle was kept in place for a minimum of 5 minutes prior to slow withdrawal at a rate of less than 0.5 mm/min. The incision was sutured with 5-0 Vicryl (VWR, 95056-936) with application of topical bacitracin, and mice were transferred to a 37°C warming pad for recovery. Animals were monitored for at least 3 consecutive days following surgery and twice a week for dramatic weight loss or any physical abnormalities until the experimental endpoint.

For virus injections, similar procedures were employed but using a 1 μL Hamilton syringe (Hamilton, 80100). For injection of AAV into the basal forebrain, the following stereotaxic coordinates were used: A/P +1.2 mm, M/L 0 mm, and D/V −5.0 mm. Other GBO transplantation sites were chosen based on experimental necessity. For retrograde monosynaptic tracing, GBOs were transplanted into either M1, S1, RSP, or ventral hippocampus (all coordinates as described above), with transplantation into multiple sites for a subset of the experiments. For Ca^2+^ imaging and electrophysiology experiments, GBOs were transplanted into the dorsal hippocampus. For anterograde tracing HSV experiments, GBOs were transplanted either in the RSP or the dorsal hippocampus. For transplantation of CHRM3 knockdown GBOs, the following stereotaxic coordinates were used: dorsal hippocampus (coordinates above) or striatum (A/P +1.0 mm, M/L +1.7 mm, D/V −3.5 mm).

### Monosynaptic viral tracing with GBOs

Transsynaptic retrograde labeling with rabies virus requires expression of helper proteins including the rabies virus glycoprotein (G) and the TVA receptor in the starter (postsynaptic) GBO population^10–12^. To enhance viral transduction efficiency, GBOs were first dissociated into a single-cell suspension with the brain tumor dissociation kit as described above. Subsequently, a retrovirus encoding dsRedExpress, rabies glycoprotein (G), and the mammalian TVA receptor (CAG-dsRedExpress-T2A-Rabies G-IRES-TVA; RTG) was incubated with 10 μg/mL polybrene (Millipore Sigma, TR1003G) on ice for 1 hour and then added to the resuspended cells. 20,000 cells together with 10 μL retrovirus in 50-100 μL culture medium were added per well in low-attachment 96-well U-bottom plates (S-bio, MS-9096UZ) to reaggregate GBOs without orbital shaking. The next day, reaggregated GBOs were transferred to ultra-low attachment 6-well culture plates (Thermo Fisher Scientific, 07-200-601) and cultured on an orbital shaker as described above. DsRedExpress signal could typically be detected via a fluorescence microscope 4-6 days post infection.

For ΔG rabies virus pre-infection retrograde tracing experiments, starter GBOs expressing RTG were incubated with EnvA-pseudotyped G-Deleted EGFP rabies virus (Salk, 32635) in low-attachment 96 well U-bottom plates. Each GBO (∼50,000 cells) was incubated with 1 μL ΔG rabies virus in 50 μL GBO culture medium per well overnight. The next day, GBOs were washed three times with DPBS and dissociated into a single-cell suspension as described above. GBO cells were resuspended and kept in ice-cold Hibernate A before intracranial injection. For pre-infection experiments, mice were sacrificed at 3, 5, or 10 dpt. For long-term GBO transplantation and virus rabies tracing, starter GBOs expressing RTG were dissociated using the same procedure. At 1 month post transplantation, a second surgery was performed to inject 1 μL ΔG rabies virus into the same location as the initial transplantation. Mice were sacrificed 10 days following rabies virus injection.

For control experiments to rule out leakage of infection-competent rabies virus (Extended Data Fig. 2f), UP-10072 GBOs expressing RTG were pre-labeled with ΔG rabies virus as described above. At either 1 day or 5 days following pre-labeling, rabies virus was extracted from GBOs and transplanted into mice for either 9 days or 5 days, respectively. Viral extraction was accomplished by first resuspending cells in distilled H_2_O. Cells were then snap frozen on dry ice for 2 minutes before thawing at room temperature, with freeze-thaw cycling repeated 5 times. Cells were then further lysed with a 26-gauge needle. Cell death and fragmentation was confirmed via trypan blue staining. 2 μL of cell lysate (from a total of 10 μL solution extracted from 9 x 10^5^ cells) was then transplanted into the RSP per mouse as described above.

For anterograde tracing experiments with monosynaptic HSV, a mixture of 400 nL AAVs (composed of a mix of 1:20 AAV2/9-ChAT-Cre-WPRE-hGHpA and Cre-dependent AAV2/8-EF1α-DIO-EGFP-2a-TK-WPRE-pA) and 600 nL H129-LSL-ΔTK-tdTomato were injected in the basal forebrain. During the same surgery, UP-10072 GBO cells were transplanted into either the RSP or the hippocampus as described above. Mice were sacrificed 10 days after the surgery and brains were harvested for immunohistology.

### Sample preparation, immunohistology, *in situ,* and confocal microscopy

To harvest brains of animals, mice were deeply anesthetized with ketamine/xylazine/acepromazine and perfused transcardially with 10 mL ice-cold DPBS followed by 10 mL of 4% paraformaldehyde (PFA). Brains were post-fixed in 4% PFA overnight at 4°C, washed with 10 mL DPBS, and transferred to 30% sucrose at 4°C for 24 hours for cryoprotection. Brains were then sectioned in the coronal plane (Leica SM 2010R) at 40 μm thickness for processing as floating sections and stored in anti-freeze medium (Bioennolife Sciences, 006799-1L) at −20°C. For each brain, every sixth section was collected into the same well of a 24-well plate. Floating mouse brain sections were washed with DPBS, incubated in DPBS with 0.3% Triton X-100 (Sigma-Aldrich, P1379) for 1 hour, and then incubated with blocking buffer (TBS with 0.1% Tween-20 (Sigma-Aldrich, T8787-50ML), 0.5% Triton X-100, 10% donkey serum (Millipore, S30), 1% BSA (Sigma-Aldrich, B6917), 22.52 mg/mL glycine (Sigma-Aldrich, 50046-50G), and 1% Mouse on Mouse Blocking Reagent (Vector Laboratories, MKB-2213-1)) for 30 minutes. Brain slices were then incubated in diluted primary antibodies in antibody buffer (TBS with 0.1% Tween-20, 0.5% Triton X-100, and 5% donkey serum) at 4°C overnight on a horizontal shaker. The following day, slices were washed 3X in TBST (TBS with 0.1% Tween-20) for 5 minutes each and incubated with secondary antibodies diluted in antibody buffer as described above for 1-2 hours at room temperature. Unless otherwise indicated, DAPI (Thermo Fisher Scientific, D1306, 1:500) was incubated with slices simultaneously during the secondary antibody incubation. Slices were washed 3X in TBST for 5 minutes each and then mounted on a glass slide (Thermo Fisher Scientific, 1518848) in mounting medium (Vector Laboratories, H-1000-10), covered with glass coverslips, and sealed with nail polish. For SST staining, brain slices underwent an additional antigen retrieval step in 1X IHC Antigen Retrieval Solution (Invitrogen, 00-4956-58) for 15 minutes at 95°C prior to blocking.

The following primary antibodies were used: Goat anti-RFP (Biorbyt, orb11618, 1:500), rabbit anti-RFP (Rockland, 600-401-379,1:500), chicken anti-GFP (Abcam, ab13970, 1:2000), goat anti-GFP (Rockland, 600-101-215, 1:500), mouse anti-Human Nuclei (Millipore, MAB1281, 1:200), mouse anti-STEM121 (Takara, Y40410, 1:250), goat anti-ChAT (Sigma-Aldrich, AB144P-200UL, 1:200), rabbit anti-VAChT (Synaptic Systems, 139103, 1:500), mouse anti-TPH2 (Thomas Scientific, AMAb91108), rabbit anti-TH (Novus Biologicals, NB300-109, 1:500), mouse anti-KI67 (BD Biosciences, 550609, 1:500), rabbit anti-KI67 (Abcam, ab16667, 1:500), rabbit anti-cleaved caspase 3 (Cell Signaling, 9661, 1:500), mouse anti-GFAP (Millipore, MAB360, 1:500), goat anti-SOX2 (Thermo Fisher, AF2018, 1:500), mouse anti-SATB2 (Abcam, Ab51502, 1:500), rat anti-CTIP2 (Abcam, Ab18465, 1:500), rabbit anti-PV (Abcam, Ab11427, 1:500), rabbit anti-SST (Thermo Fisher, PA-5-85759, 1:250), mouse anti-NeuN (Thermo Fisher Scientific, MA5-33103, 1:500), rabbit anti-Nestin (Abcam, Ab105389, 1:500), and mouse anti-EGFR (Novus Biologicals, NB200-206, 1:500). The following secondary antibodies were used: donkey anti-goat Alexa Fluor 488 (Thermo Fisher Scientific, A-11055, 1:500), donkey anti-goat Alexa Fluor 555 (Thermo Fisher Scientific, A-21432, 1:500), donkey anti-goat Alexa Fluor 647 (Thermo Fisher Scientific, A-21447, 1:500), donkey anti-rabbit Alexa Fluor 488 (Thermo Fisher Scientific, A-21206, 1:500), donkey anti-rabbit Alexa Fluor 555 (Thermo Fisher Scientific, A-31572, 1:500), donkey anti-rabbit Alexa Fluor 647 (Thermo Fisher Scientific, A-31573, 1:500), donkey anti-mouse Alexa Fluor 488 (Thermo Fisher Scientific, A-21202, 1:500), donkey anti-mouse Alexa Fluor 555 (Thermo Fisher Scientific, A-31570, 1:500), donkey anti-mouse Alexa Fluor 647 (Thermo Fisher Scientific, A-31571, 1:500), donkey anti-rat Alexa Fluor 647 (Thermo Fisher Scientific, A-48272, 1:500), and donkey anti-chicken Alexa Fluor 488 (Thermo Fisher Scientific, A-78948, 1:500). For a subset of the retrograde tracing experiments, RFP-Booster Alexa Fluor 568 (ChromoTek, rb2af568, 1:500) and GFP-Booster Alexa Fluor 488 (ChromoTek, gb2af488, 1:500) were used during the blocking step and sections were mounted immediately after blocking.

For GBO and primary GBM tissue immunohistology, tissue pieces were transferred to 1.5 mL Eppendorf tubes using wide-bore P1000 pipette tips and washed with DPBS. Tissue was then fixed in 4% PFA at 4°C overnight, triple washed in DPBS, and cryoprotected in 30% sucrose at 4°C for at least 24 hours. Tissue pieces were then transferred to a plastic cryomold (Electron Microscopy Sciences), embedded in tissue freezing medium (TFM, General Data, 1518313), and stored at −80°C. Samples were sectioned at 16 μm (Leica, CM3050S) and mounted on glass slides stored at −20°C. For immunohistology, sectioned samples were warmed to room temperature, outlined with a hydrophobic pen, and washed with DPBS for 5 minutes to remove TFM. Samples were then taken for blocking as described above for brain slice sections. For smaller GBOs (e.g., 2000-5000 cells), organoids were directly taken for blocking following the 4% PFA fixation step in low-attachment 96-well plates without the need for cryosectioning.

For *in situ* hybridization and concurrent immunostaining, cryopreserved floating mouse brain sections were mounted on (3-Aminopropyl)triethoxysilane (Sigma Aldrich, A3648) coated glass slides using PBST (PBS with 0.1% Tween-20). Slides were then processed for *in situ* hybridization using the RNAscope Multiplex Fluorescent Reagent Kit v2 (ACD, 323270) according to the manufacturer’s protocol with minor modifications. In brief, slides were washed with PBS and baked at 42°C for 30 minutes, followed by dehydration with ethanol, hydrogen peroxide treatment, and target retrieval, primary antibody incubation (chicken anti-GFP or rabbit anti-VAChT), and post-primary fixation according to the manufacturer’s protocol (MK 51-150, Rev B, Appendix D). Slides were then incubated with RNAscope Protease Plus at room temperature for 30 minutes. Next, slides were prepared for probe hybridization (ACD, UM-323100, Chapter 4) using probes for GAD1-C2 (ACD, 400951-C2), SLC17A7-C3 (ACD, 416631-C3), SLC17A6-C3 (ACD, 319171-C3), and/or GAD1-C3 (ACD, 400951-C3) and developed with TSA Vivid 570 (ACD, 323272, 1:1500) or TSA Vivid 650 (ACD, 323373, 1:1500). Following hybridization, slides were incubated with secondary antibody (donkey anti-chicken Alexa Fluor 488 and/or donkey anti-rabbit Alexa Fluor 555) with DAPI for 1 hour at room temperature before mounting slides as described above. For identification of glutamatergic neurons, we probed simultaneously for vGLUT1 (SLC17A7) and vGLUT2 (SLC17A6) with C3 probes.

Mouse brain slices, GBO (sections or whole mount), or primary tissue sections after immunohistology were imaged with a confocal microscope (Zeiss LSM 810 or Zeiss LSM 710) as *z*-stacks with either 5X, 10X, 20X, or 40X objectives. Images were pre-processed with Zen 2 software (Zeiss) for orthogonal projection and stitching and further processed with ImageJ/FIJI (v2.1.0) for exporting and quantification.

For neuron quantification after retrograde tracing, 40 μm coronal mouse brain slices consisting of every sixth slice from +2.5 mm to −4.5 mm A/P were imaged as *z*-stacks of approximately 15 μm and orthogonally projected for analysis. For each section, we annotated the number of cells in ‘level 1’ (e.g., thalamic nucleus or cortical area/layer) and ‘level 2’ regions (e.g., thalamus or isocortex) according to the coronal Allen Brain adult mouse brain atlas (http://atlas.brain-map.org)^99^ in either the ipsilateral or contralateral side to GBO or NPC injection. For each ‘level 1’ region, we identified the number of starter (GFP^+^DsRed^+^) cells and the number of input neurons (GFP^+^DsRed^-^). For starter cells in which cell number was too numerous to be quantified manually, we used an approach where we estimated the cell density (cell number per area) and then extrapolated the number of starter cells. Representative coronal sections in Fig. 2a-d were colored by GFP^+^DsRed^-^ cell proportion independently for contralateral versus ipsilateral regions. Schematic images were obtained by querying the Allen Brain Atlas API (atlas ID: eq602630314).

### Neural progenitor cell preparation and transplantation

To express the RTG helper proteins in NPCs, 200,000 cells were mixed with 100 μL of polybrene-treated retrovirus in 100 μL of F3 medium in an Eppendorf tube for 1 hour at 37°C. Cells were then seeded on six-well plates pre-coated with 1% Matrigel. After 24 hours, the medium was completely replaced with fresh F3 medium. DsRed expression could be observed under a fluorescence microscope by 5 days following retroviral infection. To pre-infect NPCs for monosynaptic tracing, 500,000 cells were infected overnight with 3 μL ΔG rabies virus in 1 mL medium in a 6-well plate. The next day, NPCs were detached with Accutase, washed 3X in DPBS, and resuspended in Hibernate-A for transplantation into the RSP as described above at 200,000 cells per mouse.

### Expansion microscopy

4.5X expansion microscopy was performed as described previously^48^ with the following modifications. In brief, mouse brain tissue sections following monosynaptic tracing experiments were pre-treated with 0.3% H_2_O_2_ in DPBS for 15 minutes at room temperature, followed by three 5-minute washes with DPBS. Immunostaining of the tissue was then performed with blocking and primary antibody incubation steps as described above (with anti-rabbit VAChT and anti-goat RFP). Tyramide-based signal amplification was performed by sequential and iterative labeling with HRP-labeled secondary antibodies (Jackson ImmunoResearch, HRP-conjugated donkey anti-goat, 703-035-147; HRP-conjugated donkey anti-rabbit, 711-035-152; all 1:500) for 60 minutes at room temperature, followed by washing 3X for 5 minutes each with TBST, then TSA labeling with either CF568 (Biotium, 10119-198) or CF660R (Biotium, 89493-550) in Tyramide Amplification Buffer Plus (Biotium, 22029) as described by the manufacturer. HRP quenching was performed in each iteration after TSA labeling by incubation with 0.3% H_2_O_2_/0.1% NaN_3_ in PBS for 15 minutes at room temperature. Next, the gelation chamber was prepared using Sigmacote (Sigma Aldrich, SL2-25mL) passivized No. 1.5 cover slips with spacers fashioned from hand-cut No. 0 glass coverslips. Imaging was performed in glass-bottomed 6-well plates with No. 0 coverslips (MatTek Life Sciences, P06G-0-20-F).

### GBO calcium imaging and analysis

To examine spontaneous Ca^2+^ transient dynamics in GBOs, organoids were seeded on top of 6-well cell culture plates containing a Millicell culture inset (0.4 μm pore size, 30 mm diameter, PICM03050). GBOs were maintained using this air-liquid interface^53,100^ (ALI) system for at least one day with 1.3 mL of GBO culture medium in each well such that GBOs were not submerged in liquid but rather open to air on one side. This allowed for the basal surface of the GBO to become relatively more flattened, making structures more amenable to live imaging in a single *z*-plane. Prior to imaging, 20 μL of 1 μM Fluo-4 AM (ThermoFisher Scientific, F14021) in GBO medium was added to each GBO at the ALI surface for at least 30 minutes. Live imaging was performed on a confocal microscope (Zeiss LSM 710) equipped with an enclosure maintaining samples at 37°C with 5% CO_2_. Each organoid was imaged at 2 Hz with the 20X objective for either 5 or 10 minutes. Following baseline recordings, 20 μL of GBO medium containing 1 mM ACh with or without a combination of the M3 muscarinic receptor antagonist 4-DAMP (100 μM) and pan-nicotinic receptor antagonist mecamylamine (100 μM, Abcam, ab120459) was added to each GBO at the ALI surface 30 minutes prior to imaging with the same conditions. Liquid at the ALI surface typically dissipated by 5 minutes after addition.

Recordings were exported as .CZI files from Zen 2 software and imported to ImageJ/FIJI for further analysis. For each organoid recording, motion artifacts were corrected with the Linear Stack Alignment with SIFT function with default parameters as necessary. Cells were then segmented by first performing a *z*-projection with ‘Average Intensity’ settings and then drawing individual ROIs for all cells in a 200 μm^2^ region for UP-10072 (and in a 300 μm^2^ region for UP-9096 and UP-9121). Intensity traces over time for individual cells were exported as .xlsx files and imported into R for analysis. To generate dF/F traces, we used the same method as above for slice Ca^2+^ imaging but with a 100-frame moving window. Traces were smoothed twice (triangular moving average) using a 5-frame window, and peaks were called using the findpeaks function in R with nups = 5, ndowns = 5, and minpeakdistance = 5.

To evaluate the immediate Ca^2+^ response of GBOs to ACh stimulation, GBOs were reaggregated into 2500 cells overnight as described above and seeded onto a Matrigel-coated 24-well plate (1:60 v/v in DMEM:F12) for at least 1 hour prior to imaging to ensure organoid adherence to the plate surface. 30 minutes before imaging, Fluo-4 AM was added to the medium to a final concentration of 1 μM. Each organoid was imaged at 2 Hz with the 20X objective on a confocal microscope as described above for 3 minutes, with ACh added to a final concentration of 1 mM at approximately the 1-minute timepoint. For antagonist experiments, either 4-DAMP or mecamylamine was added for at least 10 minutes to a final concentration of 100 μM before imaging and ACh stimulation. Similar to the Ca^2+^ transient analyses, recordings were exported from Zen 2, imported to ImageJ/FIJI, and motion artifacts were corrected as necessary. Individual ROIs were drawn, and intensity traces were analyzed in R. To generate dF/F traces, we used the same method as above for slice Ca^2+^ imaging but the baseline intensity was computed as the mean intensity of 20 frames prior to ACh stimulation. Traces were smoothed twice (triangular moving average) using a 5-frame window, and ΔdF/F_max_ was defined as the change in dF/F between the maximum and minimum intensity values across the entire trace.

### Patch-clamp recordings

For electrophysiology experiments, a mixture of 600-800 nL AAVs (composed of a mix of 1:20 AAV2/9-ChAT-Cre-WPRE-hGHpA and Cre-dependent AAV9-EF1α-DIO-hChR2(H134R)-EYFP-WPRE-hGHpA) was injected in the basal forebrain. During the same surgery, UP-10072 GBO-RTG cells were transplanted into the dorsal hippocampus. Acute slices were prepared from these animals six weeks later as previously described^101^. In brief, brains were harvested and placed immediately in ice-cold cutting solution (92 mM *N*-methyl-D-glucamine, 2.5 mM KCl, 1.2 mM NaH_2_PO_4_, 30 mM NaHCO_3_, 20 mM HEPES, 25 mM glucose, 5 mM sodium L-ascorbate, 2 mM thiourea, 3 mM sodium pyruvate, 10 mM MgSO_4_ and 0.5 mM CaCl_2_) and continuously bubbled with 95% O_2_ and 5% CO_2_. 200 μm-thick coronal sections were cut with a vibratome (Leica VT 1200S) and placed in aCSF (126 mM NaCl, 2.5 mM KCl, 1.2 mM MgSO_4_, 2.4 mM CaCl_2_, 25 mM NaHCO_3_, 1.4 mM NaH_2_PO_4_, 11 mM glucose and 0.6 mM sodium L-ascorbate) and continuously bubbled with 95% O_2_ and 5% CO_2_. Slices were incubated at 31°C for 30 minutes and then at room temperature for 30 minutes.

Brain slices were transferred into a recording chamber and perfused with oxygenated aCSF that additionally contained 200 μM 4-AP (Sigma Aldrich, A78403) and 25 μM CNQX (Tocris, 0190). DsRed^+^ GBM cells were located via a 40X water-immersion objective (Olympus BX61WI). Because of the diffusely infiltrative nature of the cells, at 6 weeks post transplantation, GBM cells could be observed in cortical, hippocampal, and subcortical regions. Recording pipettes were generated from borosilicate glass (Flaming-Brown puller, Sutter Instruments, P-97, tip resistance of 5–10 MΩ) and were filled with pipette solution consisting of 120 mM potassium gluconate, 10 mM NaCl, 1 mM CaCl_2_, 10 mM EGTA, 10 mM HEPES, 5 mM Mg-ATP, 0.5 mM Na-GTP and 10 mM phosphocreatine. Whole-cell patch clamp recordings were controlled via an EPC-10 amplifier and Pulse v8.74 (HEKA Electronik), and blue light stimulation (pE-300ultra, CoolLED, ∼25 mW) was delivered through the 40X objective. For optogenetic stimulation of cholinergic fibers, a train of 10 ms pulses at 20 Hz for 10 seconds was delivered. M3 receptor blockade was achieved by bath perfusion of aCSF consisting additionally of 100 μM 4-DAMP (Santa Cruz, sc-200167). Data were exported as raw traces in ACS file format using PulseFit (HEKA Electronik) and imported into R (v4.3.1) for analysis. For representative traces, raw traces were smoothed twice (triangular moving average) with a 25-millisecond window using the rollmean function as implemented in the zoo package in R. For analysis of maximum membrane depolarization from baseline for current-clamp traces, data were first smoothed twice using a 5-second window. Light-induced depolarization was quantified as the difference between the mean baseline voltage prior to stimulation and the maximum voltage value in the 20 seconds following the start of stimulation.

### Slice calcium imaging and analysis

UP-10072 GBOs expressing jrGECO1a were generated using lentiviruses with procedures as described above. A mixture of 600-800 nL AAVs (composed of a mix of 1:20 AAV2/9-ChAT-Cre-WPRE-hGHpA and Cre-dependent AAV9-EF1α-DIO-hChR2(H134R)-EYFP-WPRE-hGHpA) was injected in the basal forebrain. During the same surgery, UP-10072 GBO-jRGECO1a cells were transplanted into the dorsal hippocampus. At six weeks post-surgery, acute slices were prepared as described above for patch-clamp recordings. Live Ca^2+^ imaging was performed with a confocal microscope (Zeiss LSM 710) on a 10X objective by acquiring images at 2 Hz in the 555 nm wavelength channel. Light pulses for optogenetic stimulation of local cholinergic axon terminals were delivered by a laser (LRD-0470-PFFD-00100-05) with 470 nm wavelength (∼25 mW power) connected to a power supply (PSU-H-LED), with pulse length and frequency set by a programmable pulse generator (Master 8). Simultaneous Ca^2+^ imaging and optogenetic stimulation (10 ms pulses at 20 Hz for 10 seconds) were performed with or without the presence of 100 μM 4-DAMP in oxygenated aCSF. For a given slice, an inter-stimulus interval of at least 5 minutes was maintained.

Ca^2+^ imaging recordings were exported as .CZI files with Zen 2 software and imported to ImageJ/FIJI for quantification. Analysis was performed on GBM cells that consistently exhibited Ca^2+^ transients in response to optogenetic stimulation. To generate the dF/F traces, we first generated the baseline intensity trace by computing the tenth percentile of a moving 50-frame window of each raw trace using the rollapply function in R. The dF/F trace was then defined as the 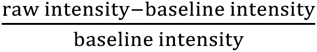 at each timepoint. The trace was then smoothed twice (triangular moving average) using a 7-frame window. We defined ΔdF/F_max_ as the maximum change in dF/F between the mean dF/F in a 10-second window prior to light stimulation compared to maximal dF/F.

### GBO migration and invasion assays

GBOs were reaggregated as described above at 2000 cells per well in a U-bottom 96 well plate for at least 24 hours prior to the assay. 6-well plates were coated with Matrigel (1:60 v/v in DMEM:F12), and 6-8 reaggregated GBOs were seeded in 2 mL of GBO culture medium per well. Treatments (to a final concentration of 1 mM ACh and/or 100 μM 4-DAMP in GBO culture medium) were added to the wells 4 hours after seeding to ensure that GBO adhesion was not affected. For experiments with CHRM3 knockdown GBOs, only ACh was added. Images were collected using an inverted phase contrast microscope (Axiovert 40 CFL) and Zen 2 software. The area covered by GBO cells after 48 hours was measured by using the ‘oval’ ROI function in ImageJ/FIJI to draw the largest bounding oval that captured the invading cells. Organoids that landed too closely to each other in the 6-well plate were excluded from analyses.

### Assembloid generation with sliced neocortical organoids and GBOs

To generate GBO-cortical organoid assembloids, we used at least 100-day old SNOs to provide a neuronal microenvironment for GBO integration. SNOs were sliced at a 300 μm thickness using a vibratome (Leica VT 1200S) at 0.1 mm/s speed and 1.2 mm vibration amplitude and transferred to ALI cultures as described above. SNOs were maintained with the ALI system with 1.3 mL of culture medium (BrainPhys Neuronal Medium (STEMCELL Technologies, 5790), 1X NeuroCult SM1 Neuronal Supplement (STEMCELL Technologies, 05711), 1X N2 Supplement (Thermo Fisher Scientific, 17502048), and 1X Penicillin-Streptomycin (Thermo Fisher Scientific, 15070063)) in each well. Medium was replenished every two days, and SNOs were cultured on ALI for at least two days prior to GBO fusion. To generate assembloids, GBOs (∼2000 – 10000 cells) were placed either directly adjacent to the SNOs such that they were physically in contact or on top of the SNOs using a P200 pipette. For invasion experiments, UP-10072 GBOs expressing RTG or expressing the shRNA construct marked by mCherry expression were used to generate assembloids to easily distinguish tumor cells from the microenvironment by combined brightfield and fluorescence imaging on a confocal microscope. All GBOs were treated with a 1-hour pulse of 1 mM ACh in GBO medium prior to assembloid generation. Images were exported from Zen 2 software and imported into ImageJ/FIJI to measure the extent of tumor invasion at the specified timepoints. The area covered by GBO cells was measured by using the ‘polygon’ ROI function in ImageJ/FIJI to draw the largest bounding polygon that captured the invading cells.

### CHRM3 shRNA knockdown analyses

GBO cells were infected with lentivirus expressing mCherry and either CHRM3 shRNA or scrambled shRNA following a reaggregation procedure as described above. Knockdown was validated by qPCR after 72 hours. RNA was first extracted using Trizol reagent (Thermo Fisher Scientific, 15-596-026) and RNA microprep kits (Zymo Research, R2062) according to manufacturer’s instructions. RNA concentration was quantified by Nanodrop (ThermoFisher Scientific, ND-2000), and 100 ng RNA was taken for reverse transcription and cDNA synthesis, which were performed using the Superscript IV First-Strand Synthesis System (ThermoFisher Scientific, 18091050) based on manufacturer’s instructions. qPCR was performed with 2 μL of cDNA, 6.25 μL of SYBR Green Master Mix (ThermoFisher Scientific, 4385612), 0.5 μL of 10 μM forward primer, 0.5 μL of 10 μM reverse primer, and 3.25 μL of nuclease-free water with the following thermocycling conditions: 95°C for 30 seconds, and 40 cycles of 95°C for 15 seconds and then 60°C for 45 seconds. Expression of CHRM3 was normalized to GAPDH based on the mean of three technical replicates by the ΔC_t_ method.

To determine the size of GBOs following CHRM3 knockdown, GBO cells were infected with CHRM3 shRNA or control scrambled shRNA lentiviruses and reaggregated as described above into organoids of 2000 cells each. After 7 days, GBOs were imaged using an inverted phase contrast microscope (Axiovert 40 CFL) and Zen 2 software, and the organoid area in a single *z*-plane was quantified using the ‘polygon’ ROI function in ImageJ/FIJI.

For quantification of KI67 proportion and cCas3 intensity following CHRM3 knockdown, GBOs were collected after 7 days for whole-mount immunostaining as described above. To quantify the percentage of KI67^+^ cells, for each organoid, the number of KI67^+^/DAPI^+^ nuclei was determined for 40-60 randomly chosen DAPI^+^ nuclei. To quantify relative levels of cCas3, for each organoid, the DAPI channel image was first used to outline the perimeter of the organoid by setting a low manual threshold combined with the Analyze Particles function with size 5000 to Infinity. Within the ROI marked by the DAPI perimeter, the mean grey value intensity for the cCas3 channel was then recorded.

For *in vivo* transplantation of CHRM3 knockdown GBOs, we transplanted equivalent numbers (∼100,000) of tumor cells into either the left or right hemispheres for each mouse, with shCHRM3 cells in the right and shScramble controls on the left, following procedures described above. Mice were sacrificed after 3 weeks, and brains were sectioned for immunostaining of mCherry as described above. To quantify relative invasion area, the ‘polygon’ tool in ImageJ/FIJI was used to draw the largest bounding polygon that captured the mCherry+ cells in either the left or right hemisphere. For each mouse, five consecutive sections (the section containing the injection sites, two sections before, and two sections after) in a single stack (composed of every 6^th^ brain section) were quantified and averaged.

### Patient survival from public databases

We queried the public GBM database GlioVis^102^ (http://gliovis.bioinfo.cnio.es) to access both gene expression and patient phenotype data for the TCGA GBM^56^ (HG-UG133A) and CGGA^57^ datasets. The CGGA dataset was filtered to only include histology consistent with GBM. A gene signature score for ACh response (either long-lasting or fast) was calculated with the GSVA^103^ package in R with method = ‘gsva’, and the score was used as an input to the function surv_cutpoint as implemented in the survminer package in R to determine the optimal cutoff for high versus low expression using maximally selected rank statistics.

### Statistics

Statistical analyses were performed in R (v4.3.1), with specific tests, sample sizes, and *p*-values indicated in the figure legends. Data are shown as mean ± s.e.m. No statistical methods were used to predetermine sample sizes. Quantifications of invasion areas and organoid immunohistology were performed blinded by two independent investigators. *P* < 0.05 was considered to be statistically significant, with levels of significance denoted as (ns): *P* ≥ 0.05, \**P* < 0.05, \*\**P* < 0.01, \*\*\**P* < 0.001.

## ACKNOWLEDGMENTS

We thank the patients and their families for the generous donations of tissue specimens; other members of the Song & Ming laboratories for discussion and suggestions for this study; Brian Temsamrit, Emma LaNoce, Angelina Angelucci, and Giana Alepa for laboratory support; Andrew Morschauser and the Penn Cytomics and Cell Sorting Shared Resource Laboratory at the University of Pennsylvania for help with single-cell sorting; and Dr. Benedikt Berninger at University College of London for providing the retroviral helper plasmid. Several schematic illustrations were created or modified from Biorender.com. This work was supported by the National Institutes of Health (R35NS116843 to H.S. and R35NS097370 to G-l.M.), Dr. Miriam and Sheldon G. Adelson Medical Research Foundation (to G-l.M., D.G., and R.K.), the Pennsylvania Department of Health (to G-l.M.), the National Science Foundation (1949735 to G.W.), the Abramson Cancer Center Glioblastoma Translational Center of Excellence (to Z.A.B. and D.M.O.), the Templeton Family Initiative in Neuro-Oncology (to Z.A.B. and D.M.O.), the Maria and Gabriele Troiano Brain Cancer Immunotherapy Fund (to Z.A.B. and D.M.O.), and the Medical Scientist Training Program at the University of Pennsylvania (to Y.S. and K.H.P.).

## AUTHOR CONTRIBUTIONS

Y.S. led the study and performed most of the analyses. X.W. and Y.S. conducted *in vivo* transplantation experiments. X.W. and Z.Z. generated the relevant viral vectors. Z.Z. generated cortical organoids used for sequencing and assembloid generation. X.W. and Y.S. performed GBO generation and culture. Y.S., X.W., and D.Y.Z. contributed to library preparation and sequencing. Q.W., D.G., and R.K. performed sequencing. Y.S., X.W., and D.Y.Z. performed immunohistology and *in situ* analyses. J.P.B., Y.W., M.M., J.G., and M.F. contributed to electrophysiology experiments. Y.S., Y.W., J.P.B., and M.M. contributed to slice Ca^2+^ imaging experiments. H.W. and F.X. provided the HSV construct. G.W. and T.G. provided the JRGECO1a construct. W.D., F.Z., K.H.P., A.S., Q.Y., S.H.K., and K.M.C. contributed to additional experiments and data collection. Z.A.B., H.I.C., E.L.P., S.S., M.P.N., and D.M.O. contributed to patient tissue collection. Y.S., X.W., G-l.M., and H.S. conceived the project, designed experiments, and wrote the manuscript with inputs from all authors.

## EXTENDED DATA FIGURES

**Extended Data Fig. 1.**
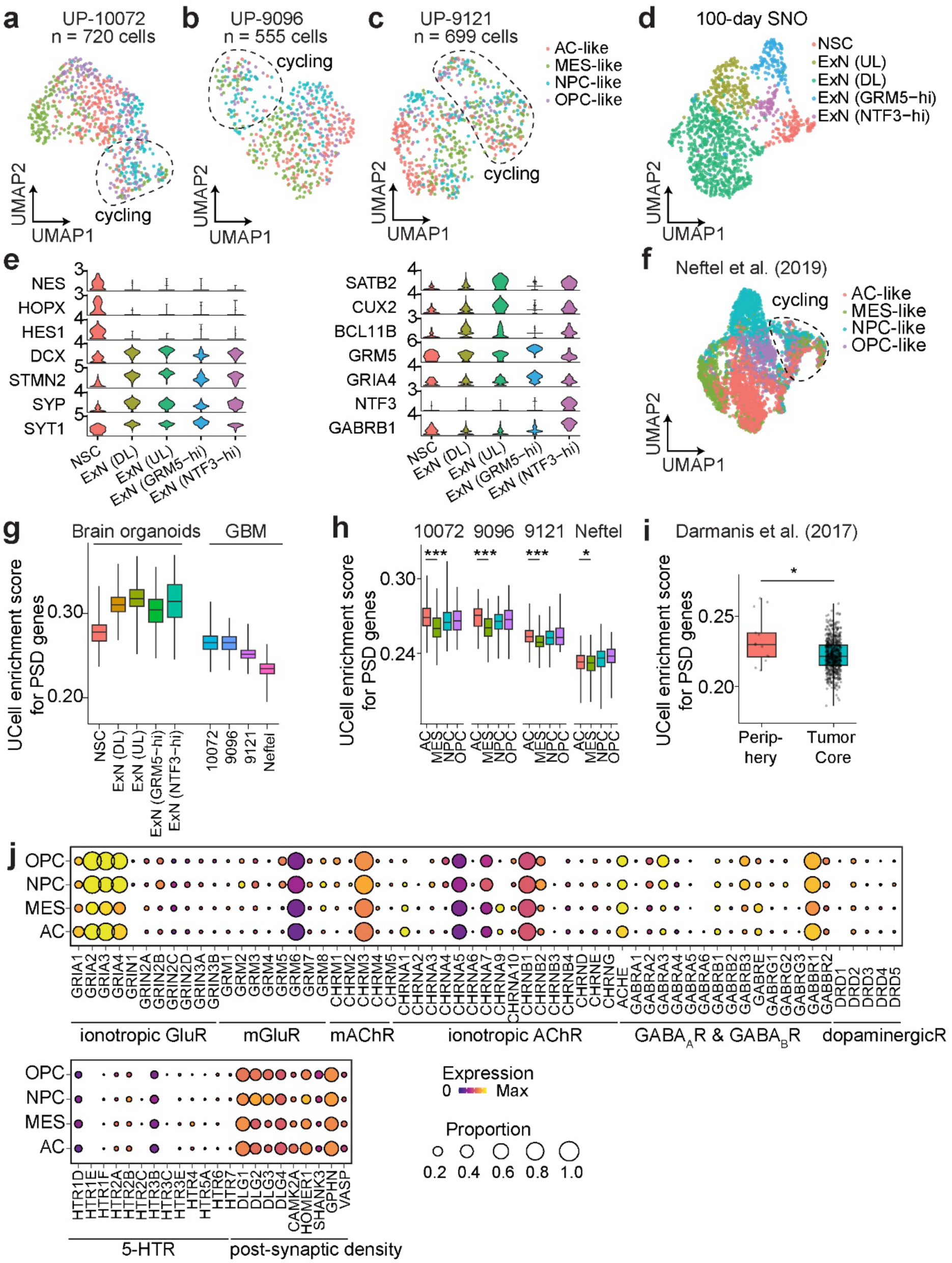
Synaptic gene enrichment in IDH-wt GBM and GBOs. **a-c**, Uniform manifold approximation and projection (UMAP) plots of scRNAseq data from UP-10072 GBOs (**a**, *n* = 720 cells across *n* = 6 organoids, mean *n* = 5525 genes per cell), UP-9096 GBOs (**b**, *n* = 555 cells across *n* = 6 organoids, mean *n* = 6153 genes per cell), and UP-9121 GBOs (**c**, *n* = 699 cells across *n* = 6 organoids, mean *n* = 6937 genes per cell). Cells are colored by assigned cell state via marker genes defined by Neftel et al^13^. Circled clusters represent cycling cells as determined by expression of *MKI67*. AC, astrocyte; MES, mesenchymal; NPC, neural progenitor cell; OPC, oligodendrocyte progenitor cell. **d**, UMAP plot of 100-day human iPSC-derived sliced neocortical organoids (*n* = 1157 cells from *n* = 3 organoids), colored by cluster identity. **e**, Violin plots of representative marker genes used to determine cell identity of cortical organoid clusters. **f**, Same as **a-c** but for Neftel et al.^13^ adult primary GBM (*n* = 4916 cells). **g**, Plot of relative enrichment score of post-synaptic density genes (GO: 0014069)^104^ across cortical organoid-derived nonmalignant clusters versus glioma cells. **h**, Same as **g** but for glioma datasets split by cell state. Comparison between AC and MES; UP-10072, *P* = 1.9 x 10^-12^; UP-9096, *P* = 5.6 x 10^-14^; UP-9121, *P* = 7.8 x 10^-12^; Neftel, *P* = 0.018; two-sided Mann-Whitney tests. **i**, Same as **g** but for enrichment by spatial location (periphery versus tumor core) with Darmanis et al.^30^ dataset (*n* = 665 cells). *P* = 0.029; two-sided Mann Whitney test. For box plots, the center line represents the median, the box edges show the 25^th^ and 75^th^ percentiles, and whiskers extend to maximum and minimum values. **j**, Gene expression dot plot as in Figure 1a, but comparing between cell states for the Neftel et al.^13^ dataset. \**P* < 0.05, \*\*\**P* < 0.001.

**Extended Data Fig. 2.**
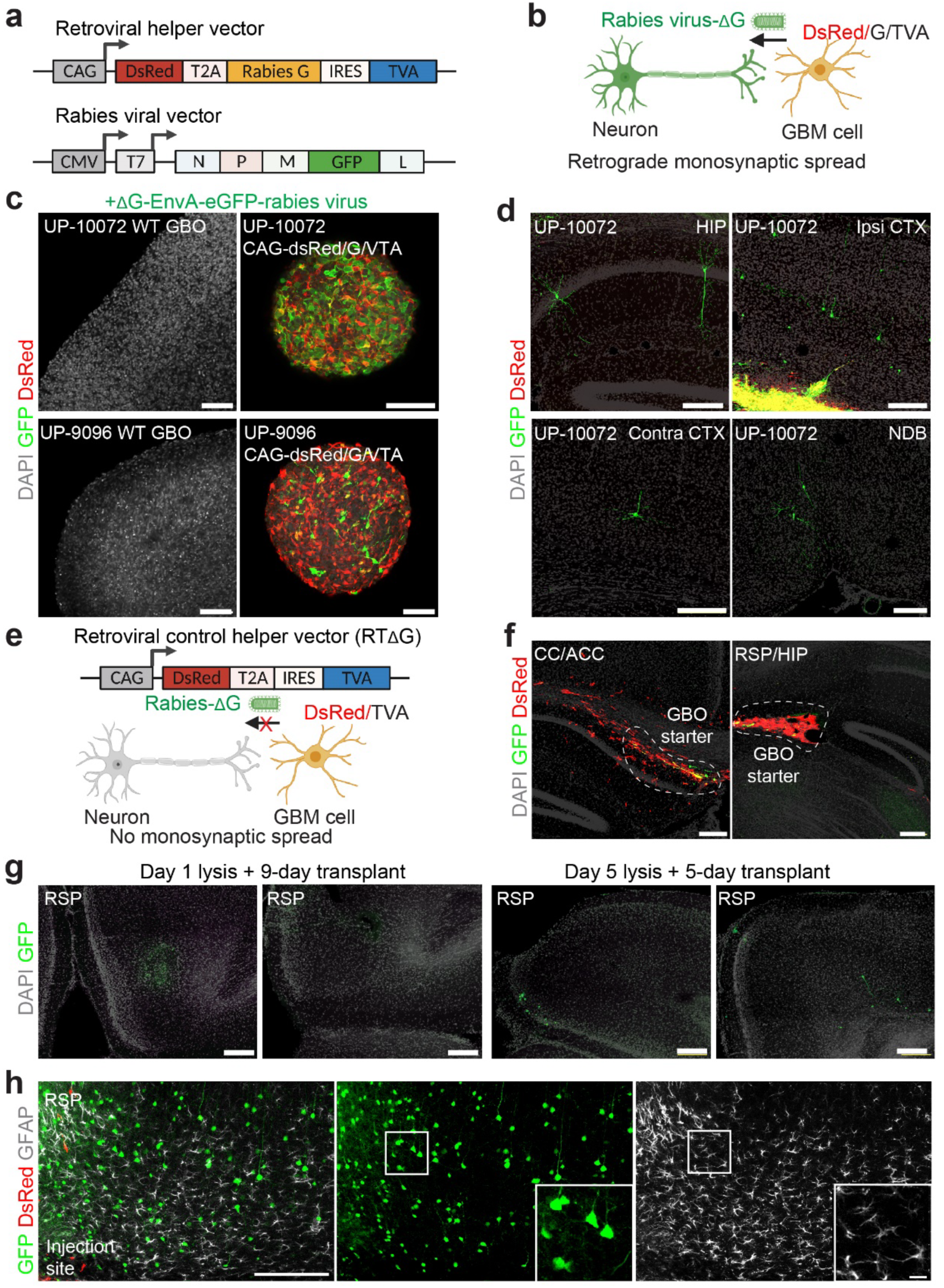
Selectivity and specificity of rabies virus transmission in GBOs. **a**, Schematic illustrations of the retroviral helper vector and EnvA-pseudotyped ΔG rabies virus vector. **b**, Principle of rabies virus monosynaptic spread. **c**, Control experiments to establish the specificity of EnvA-pseudotyped ΔG rabies towards infecting GBO cells expressing the TVA receptor protein. Nontransduced GBOs (UP-10072 or UP-9096) and GBOs expressing DsRed-G-TVA (UP-10072 or UP-9096) were infected with ΔG rabies virus for 5 days. Nontransduced (WT, wild-type) GBOs were unable to be infected by ΔG rabies virus as shown by the lack of GFP expression. Scale bars, 20 μm. **d**, Representative confocal images of monosynaptic labeling of neurons 3 dpt of pre-labeled UP-10072 cells. CTX, cortex; NDB, diagonal band nucleus. Scale bars, 200 μm. **e** – **f**, Schematic (**e**) and representative confocal images (**f**) of a control helper vector with a deletion of the G protein, thus preventing G protein-mediated transsynaptic rabies virus transmission. UP-10072 GBO cells expressing DsRed-TVA were pre-infected with ΔG rabies and transplanted into the RSP for 10 days, with no evidence of GFP^+^ neurons (*n* = 4 mice). CC, corpus callosum; ACC, anterior cingulate cortex. **g**, Representative confocal images from control experiments showing that release of rabies virus from infected GBO cells is not a significant mechanism of mouse neuronal labeling (*n* = 2 mice). UP-10072 GBOs (*n* = 3 organoids) expressing DsRed-G-TVA were pre-infected with ΔG rabies virus, lysed after 1 day and subsequently transplanted into the RSP for 9 days (Day 1 lysis + 9-day transplant). To allow for maximal viral load within GBOs prior to transplantation, the same experiment was repeated with lysis after 5 days and transplantation into the RSP for 5 days (Day 5 lysis + 5-day transplant). Low numbers of GFP^+^ mouse neurons can be observed in the latter condition, suggesting successful extraction of infection-competent rabies virus. Scale bars, 200 μm. **h**, Sample confocal immunostaining images for glial marker GFAP, showing no co-staining with GFP^+^ neurons near or far from the injection site (RSP) to establish selectivity of viral transmission. Scale bars, 200 μm (low magnification) and 20 μm (insets).

**Extended Data Fig. 3.**
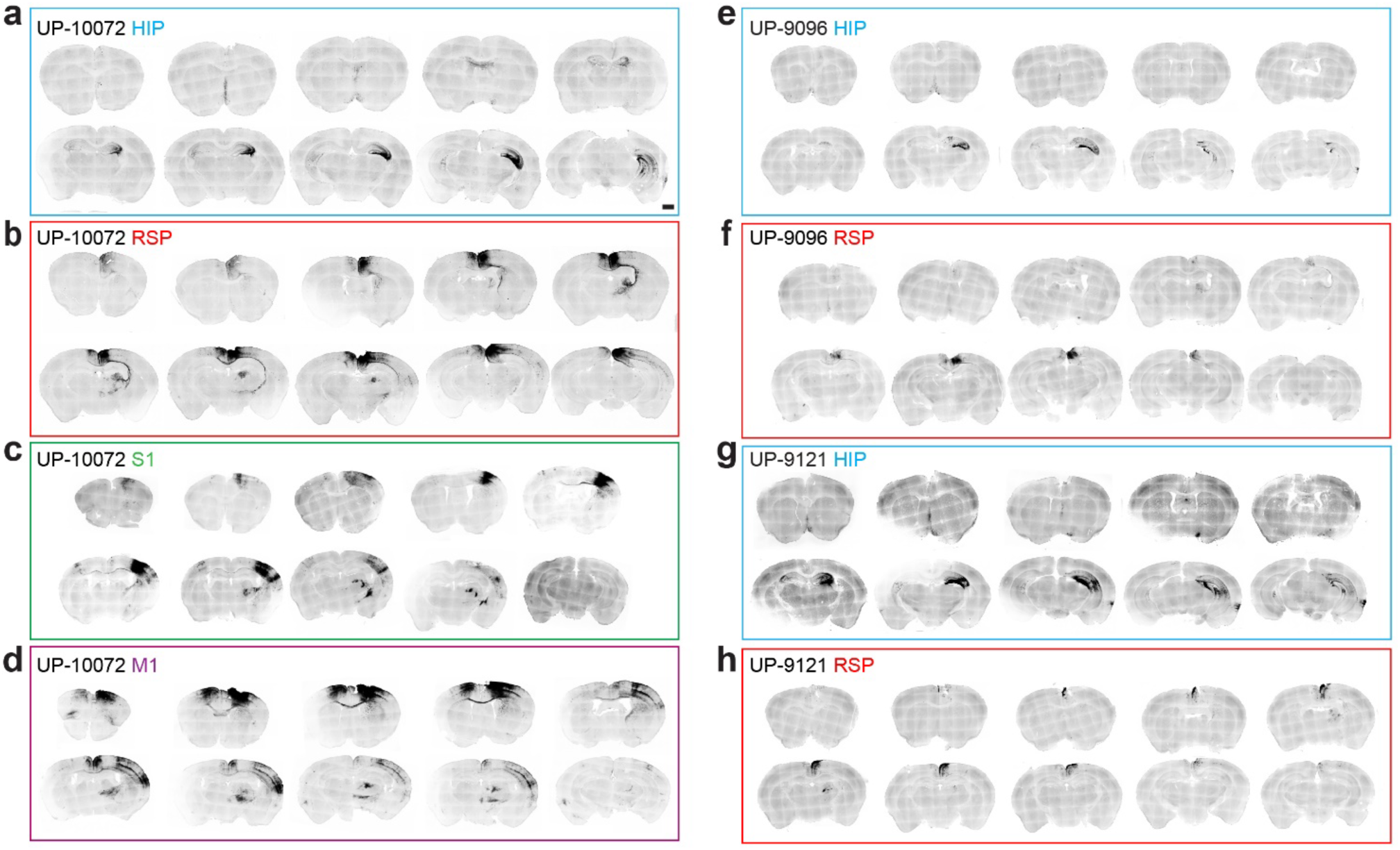
Representative brain sections across GBO transplantation sites. **a – h**, Confocal brain section images of GFP^+^ cells following transplantation of ΔG rabies virus pre-infected GBOs expressing DsRed-G-TVA for 10 days. Images are coronal sections and are arranged from anterior to posterior. Sections are arranged by GBO patients: UP-10072 (**a – d**), UP-9096 (**e, f**), UP-9121 (**g, h**). Location of transplantations include hippocampus (HIP) (**a, e, g**), retrosplenial cortex (RSP) (**b, f, h**), primary somatosensory cortex (S1) (**c**), and primary motor cortex (M1) (**d**). Scale bar, 1 mm.

**Extended Data Fig. 4.**
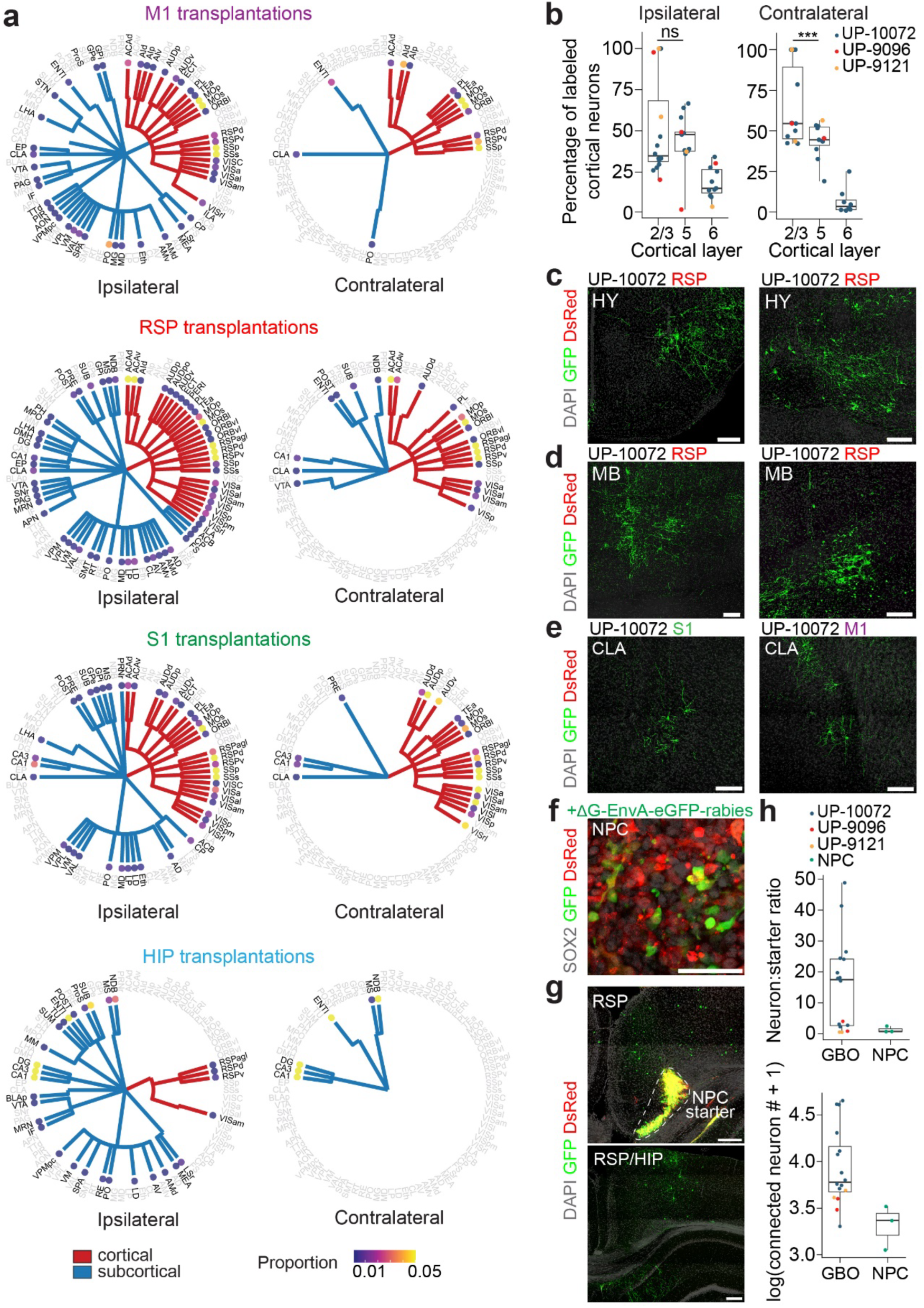
Extended characterization of monosynaptic projections onto GBM cells. **a**, Dendrogram plots showing relative proportions of input projections across cortical (red) and subcortical (blue) for ipsilateral and contralateral sites across all experiments (*n* = 16 mice as described in Figure 2e). **b**, Percentages of input cortical neurons distributed across layers for ipsilateral and contralateral neurons. Each dot represents data from one mouse (*n* = 16 mice across *n* = 3 GBOs), and data were plotted only if neurons from that cortical layer were detected. Two-tailed Student’s *t*-test, *P* = 0.44 (L2/3 vs L5, ipsilateral), *P* = 0.0082 (L2/3 vs L5, contralateral). **c – e**, Representative confocal images of GFP^+^ projections onto GBM cells from the hypothalamus (HY, **c**), midbrain (MB, **d**), and claustrum (CLA, **e**). Scale bars, 200 μm. **f**, Representative confocal images of SOX2^+^ human iPSC-derived neural progenitor cells expressing DsRed-G-TVA infected with ΔG rabies virus for 5 days. NPC, neural stem cell. Scale bar, 50 μm. **g**, Representative confocal images of monosynaptic tracing with ΔG rabies virus pre-infected NPCs transplanted into the RSP (*n* = 3 mice). Starter cells are circled (left), and images show projections either near (left) or distal (right) to the transplantation site. Scale bars, 200 μm. **h**, Comparison of neuron to starter cell ratio and total labeled neuron number for GBO transplantation (*n* = 16 mice) and NPC transplantation (*n* = 3 mice). For all box plots, the center line represents the median, the box edges show the 25^th^ and 75^th^ percentiles, and whiskers extend to maximum and minimum values. \*\*\**P* < 0.001.

**Extended Data Fig. 5.**
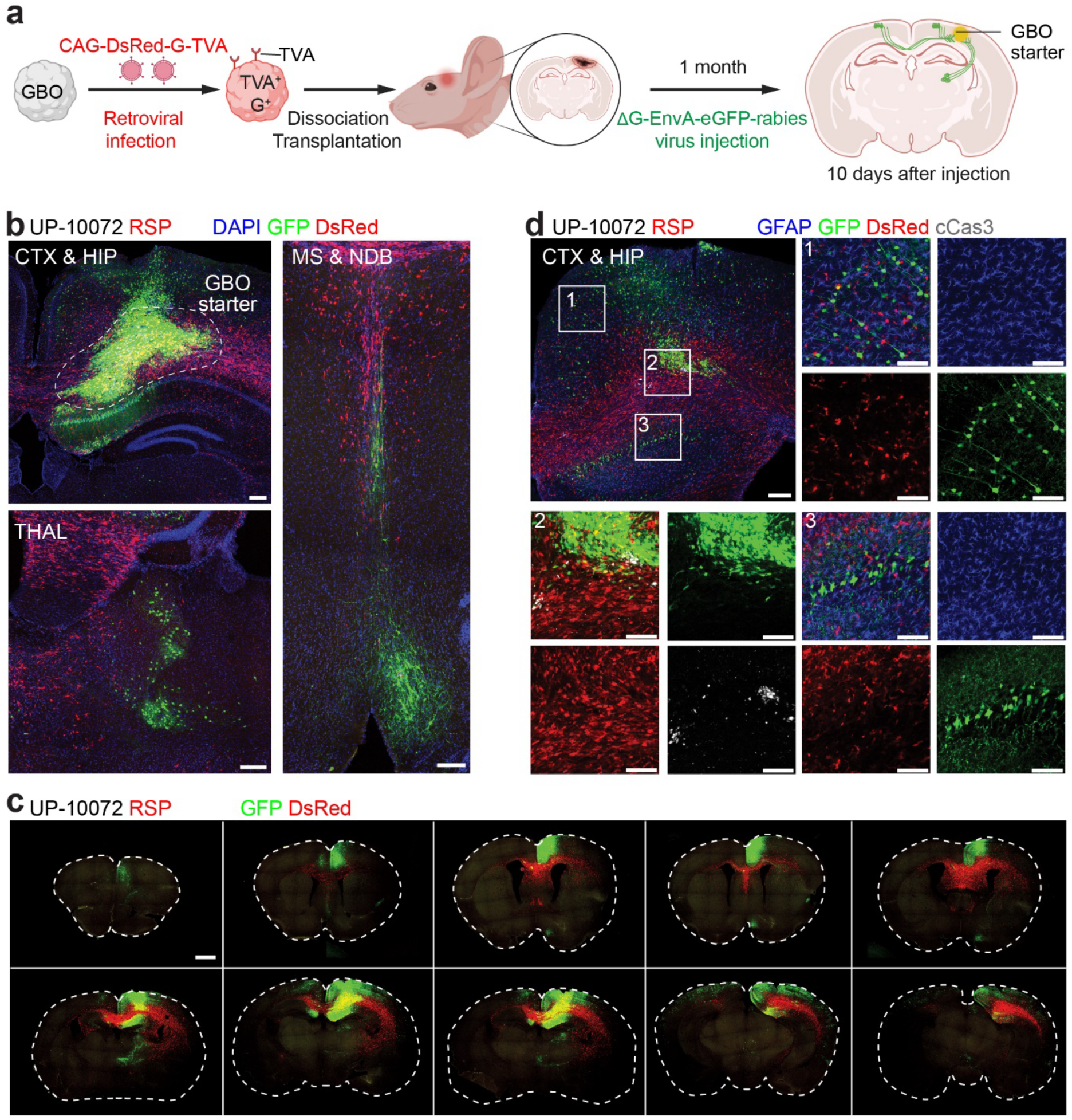
Monosynaptic tracing following long-term engraftment of GBO cells. **a**, Schematic illustration of two-step GBO retrograde monosynaptic tracing. GBOs expressing DsRed-G-TVA were transplanted into the RSP, and ΔG rabies virus was injected one month following engraftment. Mice were examined ten days following ΔG rabies virus injection (*n* = 2 mice). **b**, Representative confocal images after monosynaptic tracing, with GFP^+^DsRed^+^ GBM starter cells (circled), GFP^-^DsRed^+^ GBM cells that were unable to transmit rabies virus, and GFP^+^DsRed^-^ upstream neuronal inputs. CTX, cortex; HIP, hippocampus; THAL, thalamus; MS, medial septal nucleus; NDB, diagonal band nucleus. Scale bars, 200 μm. **c**, Representative coronal sections from anterior to posterior. Scale bar, 500 μm. **d**, Sample confocal immunostaining images for glial marker GFAP and apoptosis marker cleaved caspase 3 (cCas3), with no evidence of glial labeling by rabies virus either proximal (inset 3) or distal (inset 1) to the starter cell site and no evidence of massive cell death of GFP^+^ GBM cells at this timescale (inset 2). Scale bars, 200 μm (low magnification images) and 50 μm (high magnification images).

**Extended Data Fig. 6.**
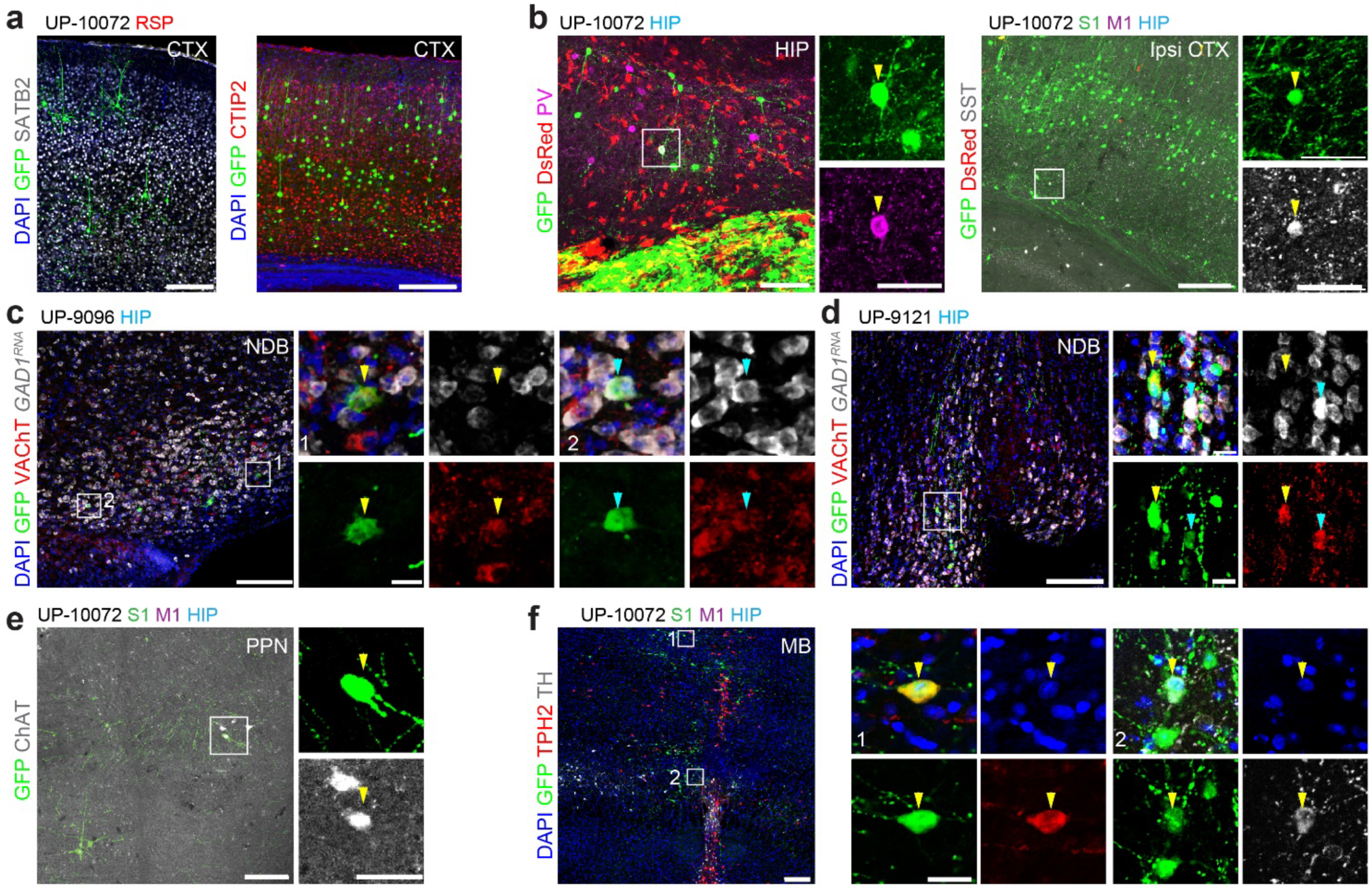
Extended characterization of input neuron identities upon GBO cell transplantation. **a**, Sample confocal immunostaining images of SATB2^+^GFP^+^ and CTIP2^+^GFP^+^ cortical glutamatergic neurons that project to GBM cells. SATB2 has a slight enrichment in upper cortical layers, whereas CTIP2 is enriched in deeper layers. CTX, cortex. **b**, Sample confocal images showing PV^+^GFP^+^ (left, hippocampus/HIP) or SST^+^GFP^+^ (right, ipsilateral cortex/ipsi CTX) GABAergic neurons. Arrowheads indicate the GABAergic neurons of interest. **c – e**, Sample confocal images of either VAChT^+^GFP^+^ or ChAT^+^GFP^+^ cholinergic neurons that project to GBM cells from either the diagonal band nucleus (NDB) (**c – d**) or pedunculopontine nucleus (PPN) (**e**). For NDB images, RNA *in situ* hybridization for *GAD1* (white) and immunostaining for VAChT and GFP were performed. Arrowheads indicate either VAChT^+^GAD1^-^GFP^+^ or VAChT^+^GAD1^+^GFP^+^ cholinergic neurons of interest. **f**, Sample confocal images showing TPH2^+^GFP^+^ serotonergic neurons (inset 1) and TH^+^GFP^+^ dopaminergic neurons (inset 2) in the midbrain (MB). For all images, GBOs and corresponding transplantation sites are as indicated. Scale bars, 200 μm (low magnification images) and 20 μm (high magnification images).

**Extended Data Fig. 7.**
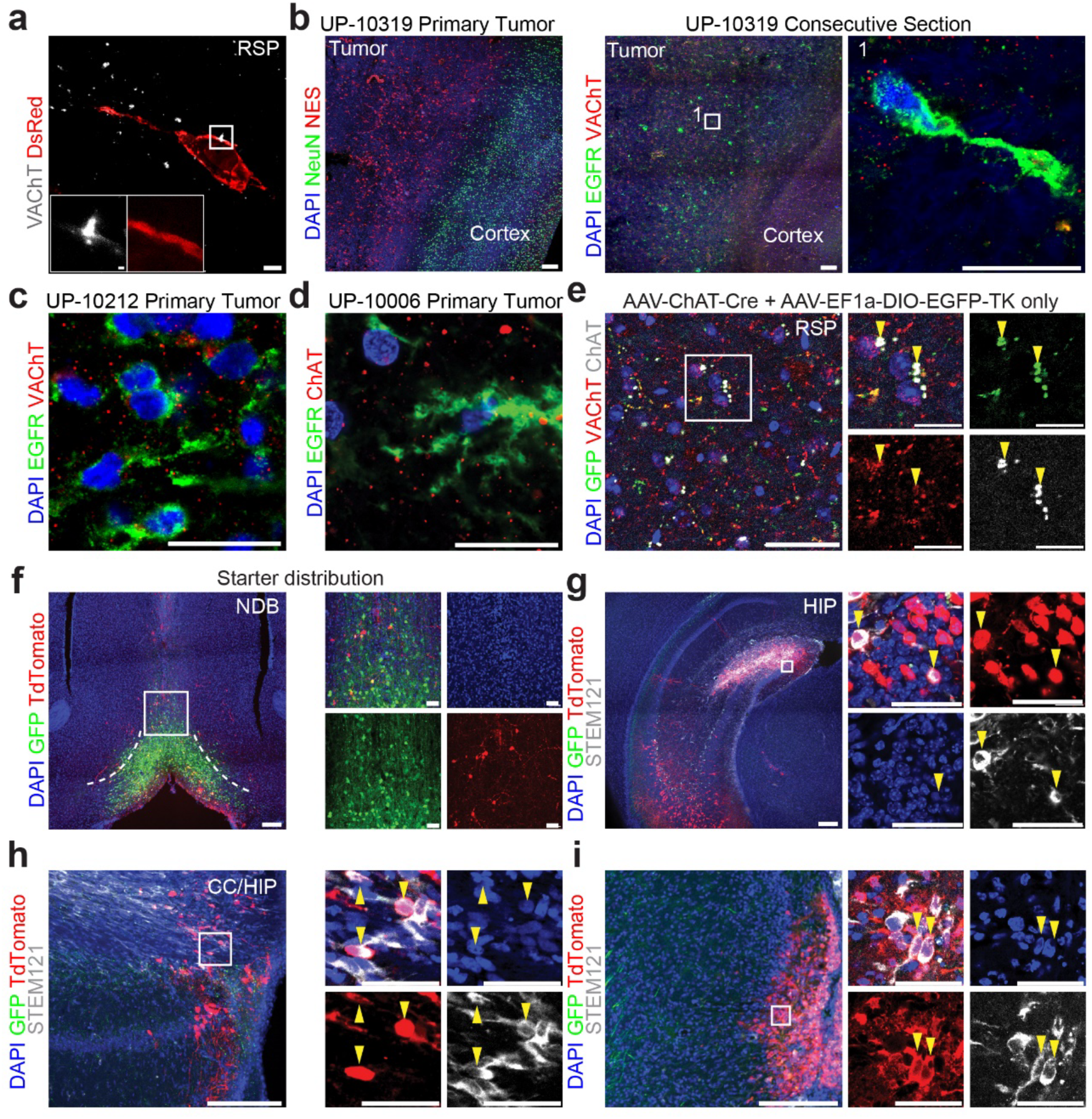
Extended validation of neuromodulatory cholinergic projection onto GBM cells. **a**, 4.5X expansion microscopy confocal image of an UP-10072 DsRed^+^ GBM cell in the RSP with VAChT^+^ puncta in close proximity. Images were taken with a 63X objective and therefore an effective 283.5X zoom. Scale bars, 4.4 μm (low magnification image) and 0.89 μm (high magnification images). **b**, Left shows confocal images of primary human GBM sample (UP-10319) immunostained for NeuN^+^ cortical neurons and NESTIN^+^ (NES^+^) tumor cells, revealing a distinct tumor-cortical boundary. Right shows confocal images of a consecutive UP-10319 section with an enrichment of VAChT^+^ puncta near EGFR^+^ tumor cells. Scale bars, 200 μm (low magnification images) and 20 μm (high magnification images). **c – d**, Confocal immunostaining images of additional primary human GBM samples (**c**, UP-10212; **d**, UP-10006) with enrichment of either VAChT^+^ (**c**) or ChAT^+^ (**d**) puncta near EGFR^+^ tumor cells. Scale bars, 20 μm. **e**, Representative confocal images of cholinergic axon terminals in the RSP (by VAChT^+^ and/or ChAT^+^ expression) 7 days after a combination of AAV-ChAT-Cre and AAV-EF1a-DIO-EGFP-TK were injected into the basal forebrain. GFP^+^ puncta co-express VAChT and/or ChAT, confirming Cre-dependent GFP expression of long-range cholinergic neurons. Arrowheads indicate examples GFP^+^ cholinergic puncta. Scale bars, 50 μm (low magnification images) and 20 μm (high magnification images). **f**, Representative confocal images of experiments to confirm monosynaptic HSV (H129-LSL-ΔTK-tdTomato) infection of starter neurons in the basal forebrain. A mixture of H129-LSL-ΔTK-tdTomato, AAV-ChAT-Cre and AAV-EF1a-DIO-EGFP-TK was injected into the basal forebrain, and immunostaining 6 days post infection revealed GFP^+^tdTomato^+^ starter cells for anterograde monosynaptic tracing. Scale bars, 200 μm (low magnification images) and 20 μm (high magnification images). **g – i**, Representative confocal images of postsynaptic GBM cells (human-specific STEM121 expression) infected by monosynaptic HSV (tdTomato expression) in the HIP (**g**), RSP (**h**), and the corpus callosum (CC)/HIP boundary (**i**). Arrowheads indicate tdTomato^+^STEM121^+^ GBM cells. Scale bars, 200 μm (low magnification images) and 20 μm (high magnification images).

**Extended Data Fig. 8.**
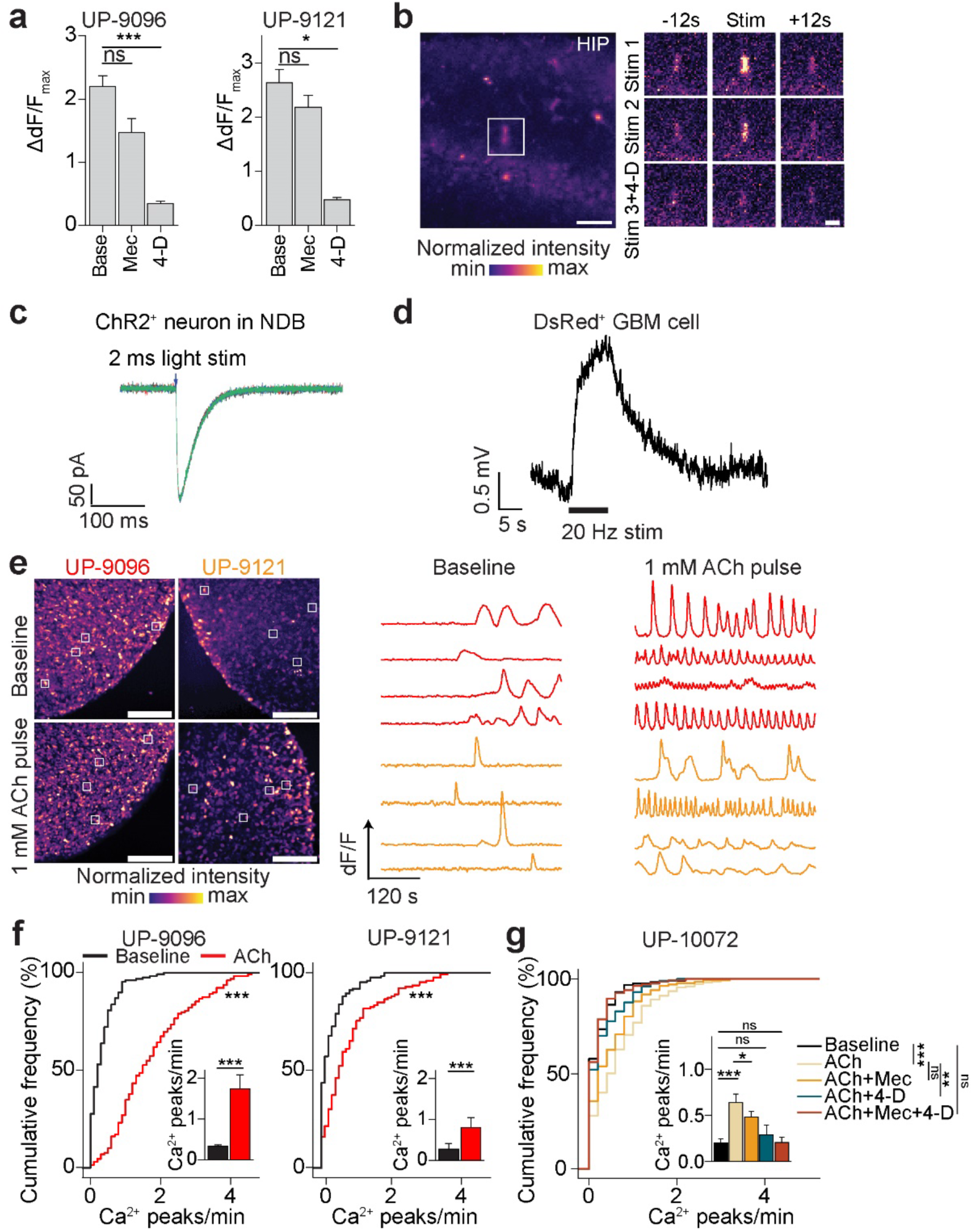
ACh-driven responses in GBM cells. **a**, Quantification of the maximal Ca^2+^ response to 1 mM ACh normalized to the baseline intensity for UP-9096 (left) and UP-9121 (right) GBOs, similar to Figure 4d. Base, baseline; Mec, 100 μM mecamylamine; 4D, 100 μM 4-DAMP. Each bar represents data from *n* = 30 cells across *n =* 3 independent GBOs. *P* = 0.068 (baseline vs. Mec, UP-9096); *P* = 0.0015 (baseline vs. 4D, UP-9096); *P* = 0.89 (baseline vs. Mec, UP-9121); *P* = 0.044 (baseline vs. 4D, UP-9121); Linear mixed-effects model fit by maximum likelihood with organoid number as a random variable; *p*-value adjustment for multiple comparisons with Tukey’s method. Data are plotted as mean ± s.e.m. **b**, Representative Ca^2+^ imaging confocal images of acute brain slices with transplanted JRGECO1a-expressing GBM cells and optogenetic stimulation of cholinergic fibers following the paradigm in Figure 4e. Inset, GBM cell of interest in the hippocampus (HIP), showing an increase in Ca^2+^ levels upon first and second stimulations but not after the addition of 100 μM 4-DAMP. Scale bars, 50 μm (low magnification image) and 20 μm (high magnification images). **c**, Inward current induced by a ChR2^+^ neuron in NDB following a 2 ms pulse of 470 nm light stimulation after acute slice preparation following the paradigm in Figure 4h, with V_m_ = −70 mV. **d**, Representative trace showing change in membrane potential of a DsRed^+^GBM cell after 470 nm light stimulation (I = 0 pA; resting membrane potential = −77 mV). **e**, Representative Ca^2+^ imaging of UP-9096 or UP-9121 GBOs at baseline or 30 minutes after a pulse of 1 mM ACh on air-liquid interface culture, similar to Figure 5a. Insets (white squares) correspond to example cells with traces shown. Scale bars, 50 μm. **f**, Cumulative distribution plots of number of spontaneous Ca^2+^ peaks per minute in UP-9096 or UP-9121 GBOs either under baseline conditions (blue) or 30 minutes after a pulse of 1 mM ACh (red), similar to Figure 5b (UP-9096: baseline, *n* = 130 cells from *n* = 3 GBOs; ACh, *n* = 129 cells from *n* = 3 GBOs; UP-9121: baseline, *n* = 124 cells from *n* = 3 GBOs; ACh, *n* = 126 cells from *n* = 3 GBOs). *P* < 2.2 x 10^-16^ (UP-9096), *P* = 4.3 x 10^-8^ (UP-9121), Kolmogorov-Smirnov test. Inset, bar plot of Ca^2+^ peaks per minute, with data plotted as mean ± s.e.m. *P* = 5.5 x 10^-33^ (UP-9096), *P* = 8.7 x 10^-10^ (UP-9121), linear mixed-effects model fit by maximum likelihood with organoid number as a random variable. **g**, Same as **f** but for UP-10072 GBOs with various receptor antagonist treatments (baseline, *n* = 133 cells from *n* = 3 GBOs; 1 mM ACh alone, *n* = 157 cells from *n* = 3 GBOs; 1 mM ACh with 100 μM Mec, *n* = 137 cells from *n* = 3 GBOs; 1 mM ACh with 100 μM 4-DAMP, *n* = 130 cells from *n* = 3 GBOs; or 1 mM ACh with both 100 μM Mec and 100 μM 4-DAMP, *n* = 137 cells from *n* = 3 GBOs). *P* = 1.35 x 10^-8^ (baseline vs. ACh), *P* = 0.12 (ACh vs. Mec), *P* = 6.15 x 10^-6^ (ACh vs 4D), *P* = 0.99 (baseline vs. Mec + 4D), Kolmogorov-Smirnov test. Inset, bar plot of calcium peaks per minute, with data plotted as mean ± s.e.m. *P* < 2.2 x 10^-16^ (baseline vs. ACh), *P* = 0.025 (ACh vs. Mec), *P* = 0.63 (baseline vs. 4D), *P* = 1.00 (baseline vs. Mec + 4D), linear mixed-effects model fit by maximum likelihood with organoid number as a random variable; *p*-value adjustment for multiple comparisons with Tukey’s method.

**Extended Data Fig. 9.**
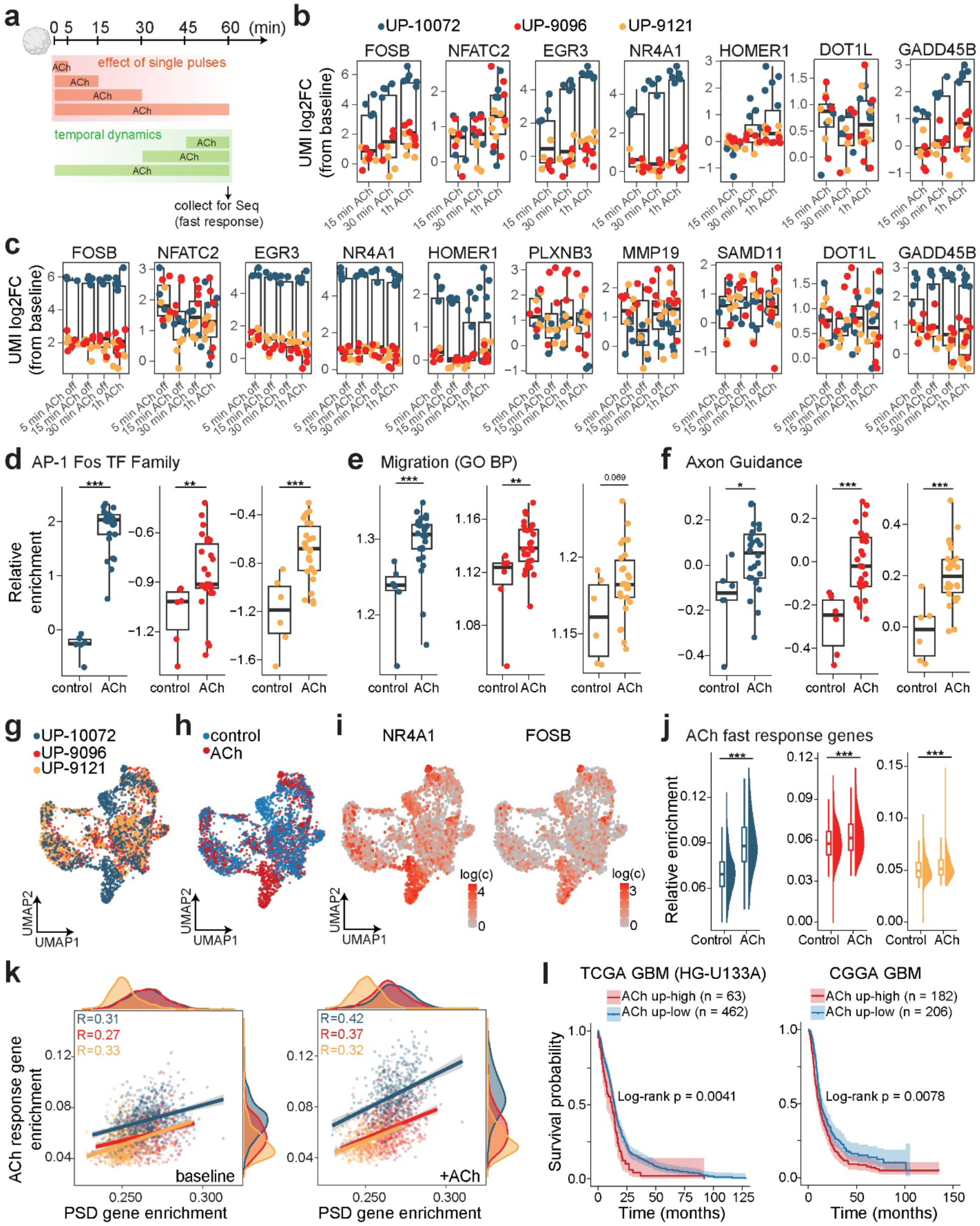
ACh-induced fast transcriptional reprogramming of GBM cells. **a**, Schematic illustration of massively parallel bulk RNA sequencing paradigm in GBOs to investigate either the transcriptional effects of short pulses of ACh or the temporal dynamics of continuous ACh treatment for various lengths of time. **b**, Plots of exemplary differentially expressed genes that vary across treatment time. Genes are plotted as log2 fold change from baseline (no ACh) values using UMI counts normalized with DeSeq2^95^. The *x* axis represents the length of time GBOs were treated with ACh prior to library preparation. Each dot represents a distinct bulk sample, and the color of the dot represents the GBO patient (*n* = 4 samples per condition per line, with a total of *n* = 3 patients). For these box plots, the center line represents the median, the box edges show the 25^th^ and 75^th^ percentiles, and whiskers extend to the ±1.5*IQR values. **c**, Same as **b** but for a set of differentially expressed genes that demonstrate long-lasting effects of a pulse of ACh. The *x* axis represents the length of time GBOs were treated with ACh, with samples taken for library preparation uniformly at 1 hour. **d – f**, Plots of module enrichment scores for the FOS transcription factor family (**d**), cell migration gene set (**e**), and axon guidance gene set (**f**) at baseline conditions or after treatment with ACh. Note that all ACh-exposed GBO samples were aggregated for the ACh condition for this analysis. Each dot represents a distinct bulk sample (*n =* 1 organoid per sample), and different GBO patients are plotted separately. AP-1: *P* < 2.22 x 10^-16^ (UP-10072), *P* = 0.0049 (UP-9096), *P* = 0.00086 (UP-9121); Migration: *P* = 0.0005 (UP-10072), *P* = 0.0071 (UP-9096), *P* = 0.069 (UP-9121); Axon guidance: *P* = 0.014 (UP-10072), *P* = 0.00066 (UP-9096), *P* = 0.00066 (UP-9121); two-sided Mann-Whitney tests. For these box plots, the center line represents the median, the box edges show the 25^th^ and 75^th^ percentiles, and whiskers extend to the maximum and minimum values. **g – h**, UMAP plots of integrated scRNAseq data of UP-10072, UP-9096, and UP-9121 (**g**, colored by patient) under either baseline or 1 mM ACh treatment conditions (**h**, colored by condition). **i**, UMAP plots of exemplary upregulated genes in response to ACh, including *NR4A1* and *FOSB*. **j**, Plots of module enrichment scores of the ACh response gene signature derived from bulk RNA sequencing experiments (**a**). *P* < 2.22 x 10^-16^ (UP-10072), *P* = 1.8 x 10^-7^ (UP-9096), *P* = 6.6 x 10^-4^ (UP-9121); two-sided Mann-Whitney tests. For these box plots, the center line represents the median, the box edges show the 25^th^ and 75^th^ percentiles, and whiskers extend to ±1.5*IQR values. Half violin plots extend to maximum and minimum values. **k**, Scatter plots of single-cell post-synaptic density (PSD) enrichment (as described in **Extended Data Figure 1g**) vs. ACh response gene enrichment in both baseline (left, no ACh) and ACh (right) conditions. Pearson correlation values are displayed and color-coded by patient. **l**, Kaplan-Meier plots of GBM patients from TCGA GBM (HG-U133A, left), and CGGA (right) datasets from GlioVis^102^. Patient profiles were grouped by GSVA score of the ACh fast response gene set, and cutoffs between high and low expression were selected using maximally selected rank statistics. Shaded areas represent the two-sided 95% confidence intervals. *P* = 0.0041 (TCGA), *P* = 0.0078 (CGGA), log-rank test. \**P* < 0.05, \*\**P* < 0.01, \*\*\**P* < 0.001.

**Extended Data Fig. 10.**
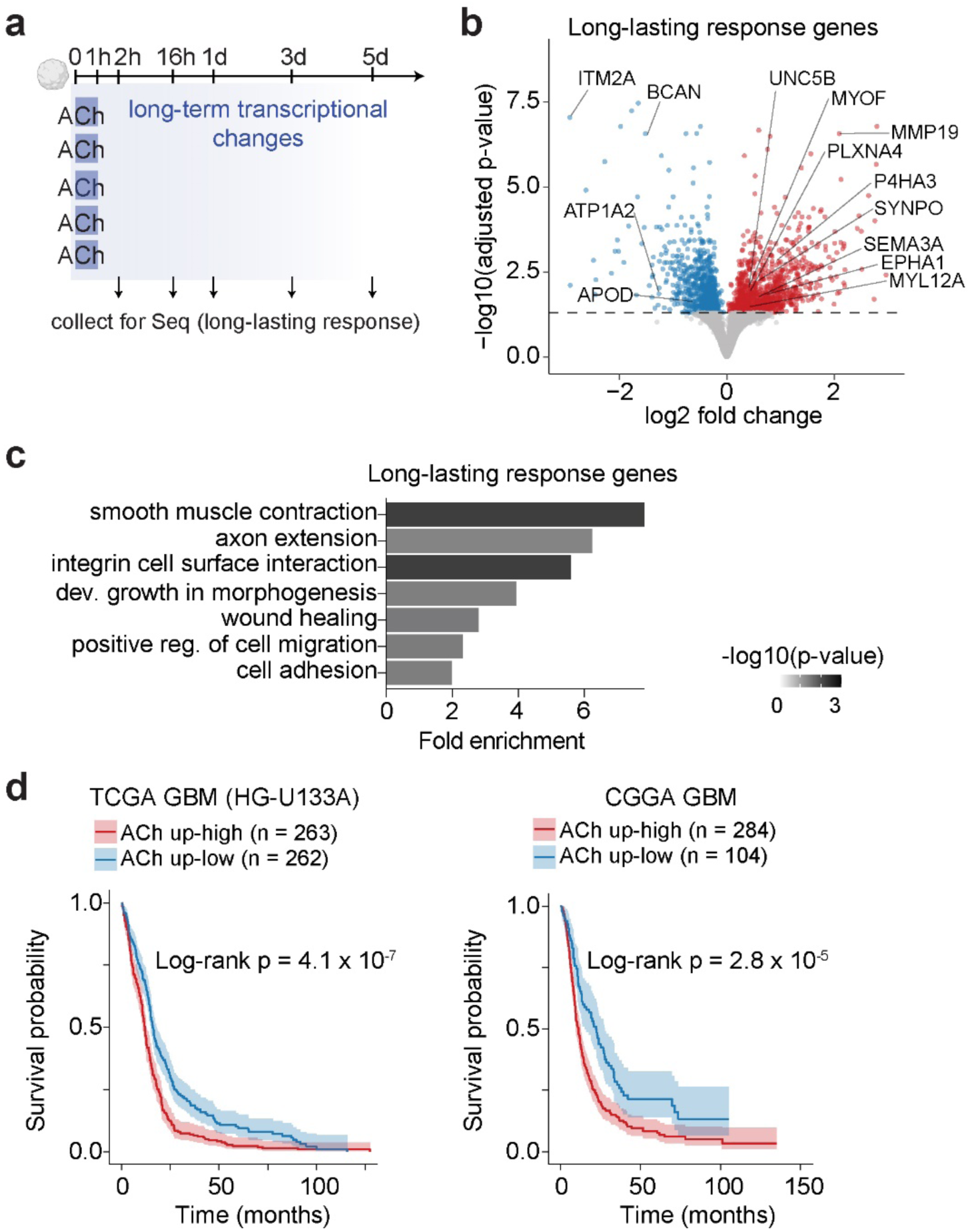
ACh-induced long-lasting transcriptional reprogramming of GBM cells. **a**, Schematic illustration of massively parallel bulk RNA sequencing paradigm in GBOs to investigate the long-term transcriptional effects of a single 1-hour pulse of ACh. **b**, Volcano plot of differentially expressed genes across UP-10072, UP-9096, and UP-9121 GBOs at 1 day following a 1-hour treatment of 1 mM ACh, defined as the long-lasting response genes. Exemplary upregulated (red) and downregulated (blue) genes are indicated. Horizontal dashed line, adjusted *p*-value cutoff of 0.05 with effect size estimation by apeglm^96^. **c**, Representative GO terms for upregulated differentially expressed genes, with the x axis indicating fold enrichment of observed genes over expected. *p-*values, Fisher’s exact test, FDR *P* < 0.05. Reg., regulation; dev., developmental. **d**, Kaplan-Meier plots of GBM patients from TCGA GBM (HG-U133A, left), and CGGA (right) datasets from GlioVis^102^. Patient profiles were grouped by GSVA score of the ACh long-lasting response gene set, and cutoffs between high and low expression were selected using maximally selected rank statistics. Shaded areas represent the two-sided 95% confidence intervals. *P* = 4.1 x 10^-7^ (TCGA), *P* = 2.8 x 10^-5^ (CGGA), log-rank test.

**Extended Data Fig. 11.**
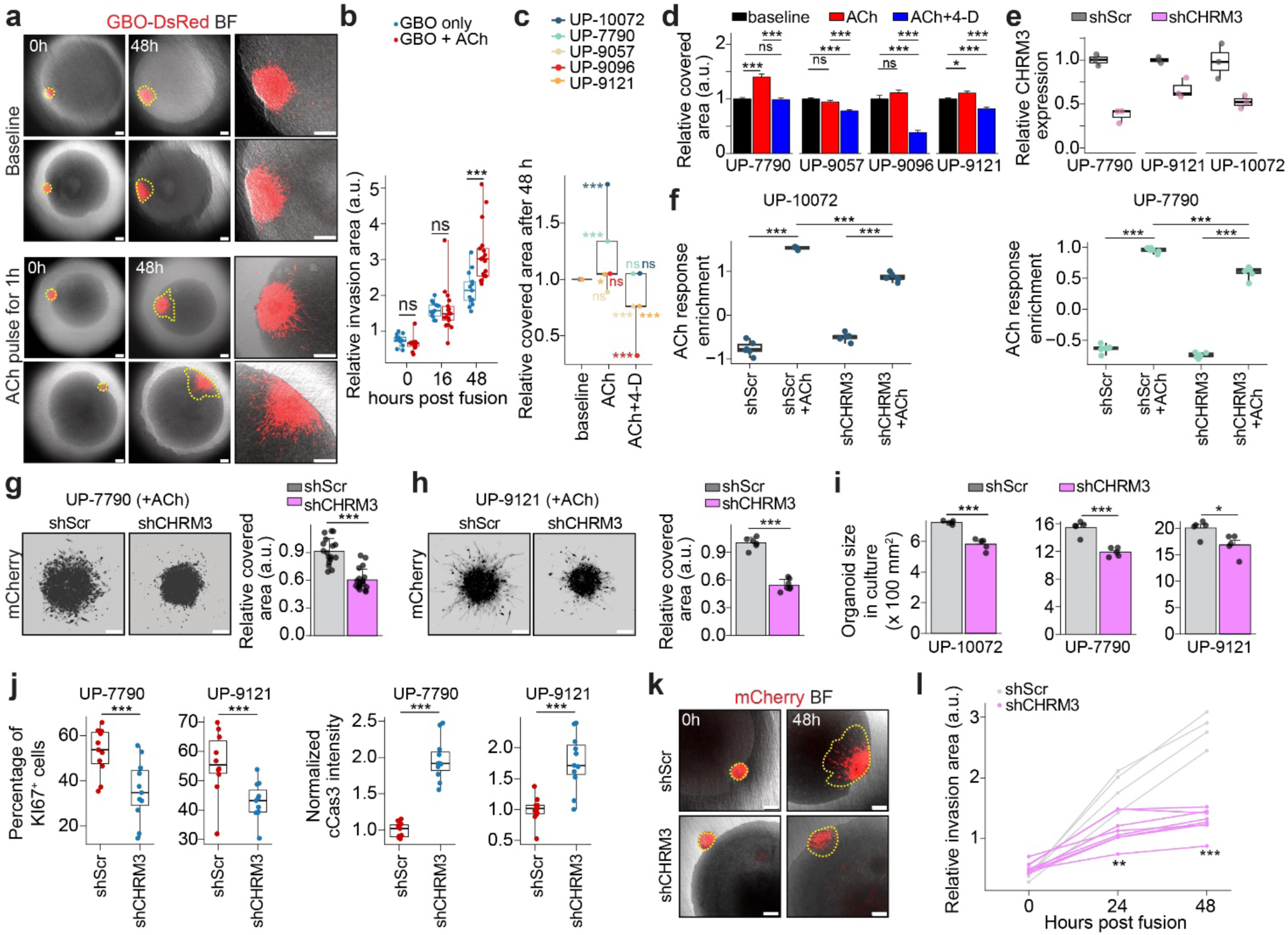
ACh-induced enhanced migration of GBM cells via CHRM3. **a**, Representative confocal images of the assembloid model with UP-10072 GBOs (red) and human iPSC-derived sliced cortical organoids (BF: brightfield). GBOs were fused with cortical organoids either under baseline conditions or after a 1-hour pulse of 1 mM ACh. Insets are zoomed in images of 48-hour timepoint. Scale bars, 200 μm. **b**, Quantification of relative invasion area of GBOs up to 48 hours post fusion. Each dot represents one individual assembloid (*n* = 13 assembloids for baseline and *n* = 17 assembloids for ACh pulse). *P* = 0.096 (0 hours), *P* = 0.902 (16 hours), *P* = 0.00062 (48 hours), two-tailed Student’s *t*-test. **c**, Quantification of migration assay data across *n* = 5 GBOs, with each color representing a different patient. Each dot represents the mean covered area by GBO cells compared to baseline for at least *n* = 12 distinct organoids per condition, which are plotted in more detail in **d**. One-way ANOVA with Tukey’s post hoc test, \**P* < 0.05, \*\*\**P* < 0.001. **d**, Quantification of Matrigel matrix-based migration assay as described above across GBOs derived from 5 patients, with *n* ≥ 12 GBOs per condition. Data are plotted as mean ± s.e.m. UP-7790: *P* = 2.8 x 10^-8^ (baseline vs. ACh), *P* = 0.95 (baseline vs. 4D), *P* = 9.8 x 10^-9^ (ACh vs. 4D); UP-9057: *P* = 0.27 (baseline vs. ACh), *P* = 1.9 x 10^-5^ (baseline vs. 4D), *P* = 9.5 x 10^-8^ (ACh vs. 4D); UP-9096: *P* = 0.30 (baseline vs. ACh), *P* = 9.6 x 10^-10^ (baseline vs. 4D), *P* = 1.1 x 10^-11^ (ACh vs. 4D); UP-9121: *P* = 0.030 (baseline vs. ACh), *P* = 2.2 x 10^-4^ (baseline vs. 4D), *P* = 1.1 x 10^-8^ (ACh vs. 4D); UP-10072: *P* < 2.2 x 10^-16^ (baseline vs. ACh), *P* = 0.14 (baseline vs. 4D), *P* = 4.0 x 10^-12^ (ACh vs. 4D); One-way ANOVA with Tukey’s post hoc test. **e**, Relative efficacy of CHRM3 knockdown via shRNA normalized to GAPDH expression assayed by qPCR in UP-7790, UP-9121, and UP-10072. Each dot represents RNA extracted from organoids from an independent lentiviral transduction (*n* = 3 biological replicates). **f**, Plots of module enrichment scores of the top ACh response genes as defined by bulk RNA sequencing of shScramble and shCHRM3 organoids for UP-10072 and UP-7790. Each dot represents a bulk RNA sequencing sample (*n* = 1 organoid per sample) under either baseline conditions or after a 1-hour pulse of 1 mM ACh. UP-10072: *P* = 1.34 x 10^-7^ (Scr vs. Scr + ACh), *P* = 6.2 x 10^-10^ (KD vs. KD + ACh), *P* = 1.1 x 10^-6^ (Scr + ACh vs. KD + ACh); UP-7790: *P* = 6.1 x 10^-11^ (Scr vs. Scr + ACh), *P* = 2.3 x 10^-8^ (KD vs. KD + ACh), *P* = 1.1 x 10^-4^ (Scr + ACh vs. KD + ACh); multiple Student’s *t*-tests with FDR correction. **g**, Representative confocal images (left) and migration area quantification (right) of UP-7790 GBOs with CHRM3 knockdown versus scrambled shRNA (*n* ≥ 17 organoids per condition), similar to Figure 5i. Data are plotted as mean ± s.e.m. Two-tailed Student’s *t-*test, *P* = 2.5 x 10^-8^. Scale bars, 200 μm. **h**, Same as **g** but for UP-9121 (at least *n* = 6 organoids per condition). Data are plotted as mean ± s.e.m. Two-tailed Student’s *t-*test, *P* = 4.2*10^-8^. Scale bars, 200 μm. **i**, Quantification of organoid size in culture 7 days following shRNA infection (*n* = 5 organoids per condition, *n* = 3 patients). *P* = 4.1 x 10^-5^ (UP-10072), *P* = 1.7 x 10^-4^ (UP-7790), P = 0.021 (UP-9121), two-tailed Student’s *t*-tests. **j**, Quantification of the percentages of KI67^+^ cells and normalized cCas3 intensity in shCHRM3 vs. shScramble GBOs (*n* ≥ 10 GBOs per patient per condition), similar to Figure 5k-l. *P* = 1.4 x 10^-9^ (UP-7790, cCas3 intensity), *P* = 4.8 x 10^-5^ (UP-9121, cCas3 intensity), *P* = 0.0026 (UP-7790, KI67 proportion), *P* = 0.0036 (UP-9121, KI67 proportion), two-tailed Student’s *t*-tests. **k**, Representative confocal images of assembloid model with either shCHRM3 or shScramble UP-10072 GBOs (red) and human iPSC-derived sliced cortical organoids (brightfield) at 0 or 48 hours post-fusion. GBOs were treated with a pulse of 1 mM ACh for 1 hour and washed prior to assembloid generation. Scale bars, 200 μm. **l**, Quantification of relative invasion area of shCHRM3 (*n* = 8 organoids) or shScramble (*n* = 4 organoids) over time. *P* = 0.0019 (24 hours), *P* = 6.2 x 10^-7^ (48 hours), two-tailed Student’s *t*-tests. For box plots in **d, e, i**, the center line represents the median, the box edges show the 25^th^ and 75^th^ percentiles, and whiskers extend to maximum and minimum values. \**P* < 0.05, \*\**P* < 0.01, \*\*\**P* < 0.001.

